# ISX-9 manipulates endocrine progenitor fate revealing conserved intestinal lineages in mouse and human

**DOI:** 10.1101/787788

**Authors:** Anastasia Tsakmaki, Patricia Fonseca Pedro, Polychronis Pavlidis, Bu’Hussain Hayee, Gavin A Bewick

## Abstract

Enteroendocrine cells (EECs) survey the gut luminal environment and co-ordinate hormonal, immune and neuronal responses to it. They exhibit well characterised physiological roles ranging from the control of local gut function to whole body metabolism, but little is known regarding the regulatory networks controlling their differentiation, especially in human gut.

The small molecule Isoxazole-9 (ISX-9) stimulates neuronal and pancreatic beta-cell differentiation, both closely related to EEC differentiation. We used ISX-9 as a tool to explore EEC specification in mouse and human intestinal organoids. ISX-9 increased the number of neurogenin3 (*Ngn3*) positive endocrine progenitor cells and upregulated *NeuroD1* and *Pax4*, transcription factors which play roles in mouse EEC specification. Single cell analysis revealed induction of *Pax4* expression in a developmentally late *Ngn3^+^* population of cells and potentiation of genes associated with progenitors biased towards serotonin-producing enterochromaffin (EC) cells. This coincided with enrichment of organoids with functional EC cells which was partly dependent on stimulation of calcium signalling in a population of cells residing outside the crypt base. Inducible *Pax4* overexpression, in ileal organoids, uncovered its importance as a component of early human endocrine specification and highlighted the potential existence of two major endocrine lineages, the early appearing enterochromaffin lineage and the later developing peptidergic lineage which contains classical gut hormone cell types.

Our data provide proof-of-concept for the controlled manipulation of specific endocrine lineages with small molecules, whilst also shedding new light on human EEC differentiation and its similarity to mouse. Given their diverse roles, understanding endocrine lineage plasticity and its control could have multiple therapeutic implications.

## Introduction

The intestinal epithelium is a key interface with our external environment. It renews itself every 4-5 days and is composed of 5 terminally differentiated cell types; the absorptive enterocytes and the secretory Paneth, goblet, tuft and enteroendocrine cells (Gehart and Clevers, 2019). These cells are constitutively generated by cycling Lgr5^+^ crypt stem cells and together they orchestrate the epithelium’s major functions, nutrient absorption and defence. Despite representing only 1% of the epithelium, the enteroendocrine cells (EECs) constitute the largest hormone producing tissue and have been described as the gut’s sentinels. They sample the luminal, circulating and local tissue environments and co-ordinate an appropriate response from the epithelium, immune and nervous systems (Gribble and Reimann, 2016). For example, they play a key role in controlling the response to a meal, fine tuning physiology to ensure optimal fuel absorption, use and storage (Sam et al., 2012). Gut hormones exhibit actions ranging from the local control of gut motility, absorption and secretion, to the regulation of whole-body metabolism (Melvin et al., 2016). Whilst there is a large body of evidence describing the functional roles of gut hormones comparatively little is known about the factors which control EEC differentiation and assign subset identity.

EECs were originally classified by immunostaining according to their dominant hormone product; Glucagon-like peptide 1 (GLP-1), Glucagon-like peptide 2 (GLP-2) and Peptide YY (PYY) are secreted by L cells, Glucose-dependent insulinotrophic polypeptide (GIP) by K cells, Somatostatin (SST) by D cells, Cholecystokinin (CCK) by I cells, Secretin (SCT) by S cells, Gastrin (GAST) by G cells, Serotonin by enterochromaffin (EC) cells, and Neurotensin (NTS) by N cells. Recent use of fluorescent reporter mice and transcriptomics has revealed EEC subsets maybe less well defined (Adriaenssens et al., 2015; Engelstoft et al., 2015; Habib et al., 2012; Knudsen et al., 2015). The hormonal repertoire of an EEC is a function of its differentiation trajectory, its location within the gut and its height in the crypt-villus axis, which dictates differential exposure to Wnt and BMP signalling gradients (Basak et al., 2017; Beumer et al., 2018). Mouse lineage tracing studies have identified a handful of transcription factors (TFs) regulating EEC differentiation. Cells exiting the stem cell compartment are fated to be secretory cells by *Notch* inhibition, followed by *Atoh1* expression (Zecchini et al., 2005) (Fre et al., 2005; Stanger et al., 2005). Atoh^+^ cells are then designated to the endocrine lineage by expression of the bHLH TF neurogenin3 (*Ngn3*) (Li et al., 2011). In mouse, TFs downstream of *Ngn3* known to be necessary for subset specification include *Insm1* (substance P, NTS) (Gradwohl et al., 2000), neurogenic differentiation 1 (*NeuroD1)* (SCT, CCK) (Mutoh et al., 1997; Mutoh et al., 1998; Naya et al., 1997), *Nkx2.2* (CCK, GAST, GIP and SST) (Desai et al., 2008), *Pax4* (5-HT, SCT, GIP, PYY, CCK) (Beucher et al., 2012) *Pax6*, *Foxa1* and *Foxa2* (Preproglucagon and its products GLP-1 and 2) (Ye and Kaestner, 2009), *Arx* (GLP-1, GIP, CCK, SCT, GAST and GHRL) (Beucher et al., 2012), and *Lmx1A* (5-HT) (Gross et al., 2016). Nevertheless, the regulatory networks controlling EEC identity have remained largely unknown, until a recent sophisticated study described a time resolved transcriptional road map of mouse EEC fate trajectories (Gehart et al., 2019). It now appears classical TFs are more promiscuous than lineage tracing implied. Furthermore, there is a paucity of knowledge regarding EEC specification in human intestinal epithelium due to lack of tractable model systems, although, several of the classical TFs are upregulated in response to a *NGN3* pulse in intestinal organoids derived from human pluripotent stem cells (Sinagoga et al., 2018; Spence et al., 2011). Understanding the factors which control gut endocrine pedigree has implications for several clinical conditions including diabetes, obesity, gut inflammatory disorders and perhaps cognitive disorders including depression and anxiety. Deciphering how to manipulate EECs may open novel treatment avenues and offer a clearer understanding of epithelial homeostasis.

To identify a candidate molecule which might influence EEC fate we drew parallels from other endocrine tissues. Gut endocrine specification is strikingly like that in the pancreas and both bear close resemblance to neuronal differentiation. The small molecule isoxazole-9, (*N*-cyclopropyl-5-(thiophen-2-yl)isoxazole-3-carboxamide (ISX-9), was uncovered in a chemical screen for drivers of neuronal differentiation (Schneider et al., 2008). It activates *NeuroD1* and has also been used to investigate pancreatic beta-cell differentiation (Dioum et al., 2011; Kalwat et al., 2016). We explored the effects of ISX-9 on EEC identity in organoids derived from mouse and human tissue resident stem cells. Our data demonstrate proof-of-concept that specific EEC populations can be manipulated with a small molecule, highlight the similarities between mouse and human EEC differentiation and provide a tool to study human enterochromaffin cells *in vitro.*

## Results

### ISX-9 increases the expression of transcription factors associated with EEC differentiation

ISX-9, increases expression of *NeuroD1* and induces differentiation of neuronal (Schneider et al., 2008), cardiac (Sadek et al., 2008) and islet endocrine progenitors (Dioum et al., 2011). We wondered if these properties could be harnessed to manipulate gut endocrine differentiation. Mouse small intestinal organoids exposed to increasing doses of ISX-9 (48-hr treatment) had increased expression of transcription factors known to be important for endocrine specification (Fig. 1A-F). As expected, ISX-9 dose-dependently increased *NeuroD1*, mimicking its role in other tissues (Fig. 1A). Interestingly, *Ngn3*, a transcription factor usually associated with the earliest identifiable endocrine progenitor cell (Li et al., 2011) and thought of being upstream of *NeuroD1*, was also increased (Fig. 1B). Other downstream TFs were differentially affected by ISX-9, with *Pax4* being increased whilst *Arx* was unaffected (Fig. 1C and D). *Atoh1* expression was inhibited at 40 and 80 μM but there was little effect on *Hes1* expression at any dose, an indirect measure of Notch signalling (Fig. 1E and F). Notch and *Atoh1* control the gate between the secretory and absorptive lineages (Li et al., 2011). These data suggest ISX-9 may act downstream of this node and could favour the differentiation of specific EEC subsets based on its opposing effects on *Pax4* and *Arx*.

**Figure 1:**
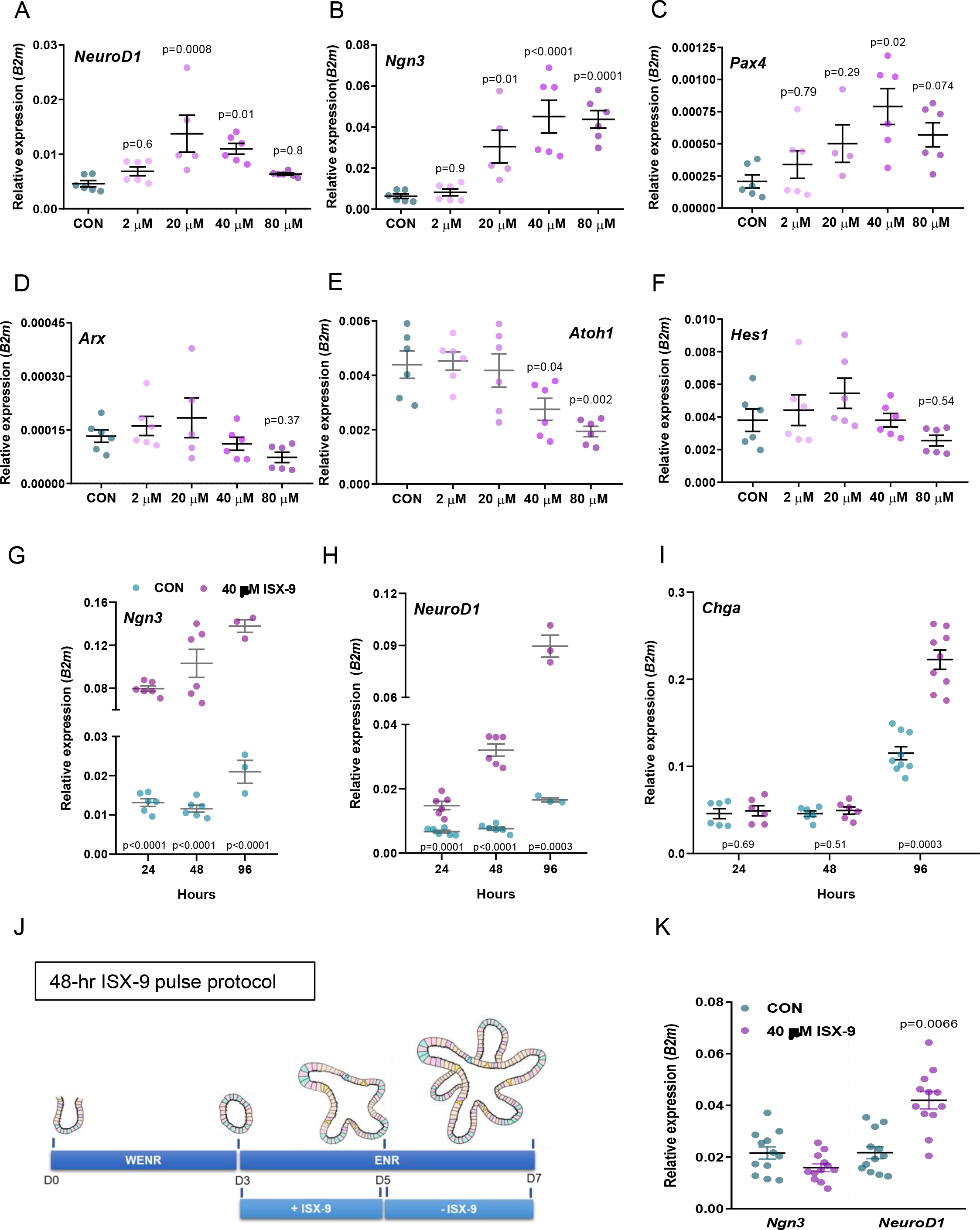
Effects of ISX-9 on the expression of transcription factors associated with EEC differentiation. (A-F) Expression of *NeuroD1* (A), *Ngn3* (B), *Pax4* (C), *Arx* (D), *Atoh1* (E) and *Hes1* (F) in mouse intestinal organoids cultured in the presence of increasing doses of ISX-9 (2 μM-80 μM) for 48 hrs. (G-I) Expression of *Ngn3* (G), *NeuroD1* (H) and *ChgA* (I) in mouse organoids after 24 hrs, 48 hrs and 96 hrs continuous exposure to 40 μM ISX-9. (J) Schematic diagram explaining our experimental paradigm (K) Expression of *Ngn3* and *NeuroD1* in mouse intestinal organoids following our experimental paradigm. Data are represented as mean ± SEM. One-way ANOVA with Dunnett post hoc test (A-F), unpaired two-tailed Student’s t test (G-I, K).

We chose to use the 40 μM dose in further experiments since at higher doses ISX-9 strongly inhibited *Atoh1* and did not significantly increase *NeuroD1*. To explore the effect of ISX-9 on gut epithelial homeostasis and EEC specification we designed the following protocol. After passage, intestinal organoid cultures were maintained in stem cell growth media (WENR) for 3 days to create a stem cell enriched baseline. We then switched to differentiation media (ENR) and exposed cultures to ISX-9 for up to 4 days, the previously reported timeframe for maturation of EECs in intestinal organoids (Petersen et al., 2014). ISX-9 increased the expression of *Ngn3* and *NeuroD1* at 24, 48 and 96 hrs but only increased chromogranin A *(Chga)*, a gene selectively expressed in terminal differentiated EECs (Engelstoft et al., 2015), at 96 hrs (Fig. 1G-I). This was highly reflective of the known differentiation trajectory of the EEC lineage. We next refined our paradigm to consist of a 48-hr pulse of ISX-9 at the beginning of the 4-day differentiation period, on day 3 of the protocol (Fig. 1J). This removed a direct effect of ISX-9 as a confounding factor and ensured measurements made at the end of the protocol could be more easily attributed to altered cell fate. In this paradigm *Ngn3* expression was unchanged whilst *NeuroD1*, which is expressed in all early to late EEC subsets, remained elevated at 96 hrs indicating the ISX-9 pulse increased EEC specification in mouse intestinal organoids (Fig. 1K).

### ISX-9 specifically enriches markers of EC cells and does not affect organoid growth

Having established a paradigm in which ISX-9 appeared to increase EEC differentiation, there was a need to determine if this was authentic manipulation of cell fate or a consequence of increasing general organoid growth. We measured the morphological characteristics of organoids at different time points during our protocol; day 3 at baseline, day 5 immediately following the ISX-9 pulse, and day 7, the end of the differentiation period. There were no obvious differences between ISX-9-treated and control organoids in their general appearance or growth rates (Fig. 2A). Equally, we found no difference between treatment and control in either surface area or perimeter of the organoids (Fig. 2B and C). We noted a trend for a reduction in the proportion of very budded ISX-9-treated organoids (those with greater than 5 buds) at day 7 (Fig. 2D). This could suggest a reduction in stem cell proliferative capacity. However, in accordance with the surface area and perimeter observations, the number of EdU^+^ cells (a marker of cells in S-phase and therefore undergoing proliferation) did not differ between treatments immediately post ISX-9 treatment or at the end of the study (Fig. 2E and F). These results expand our evidence that ISX-9 alters cell fate and increases the EEC lineage independently of organoid growth.

**Figure 2:**
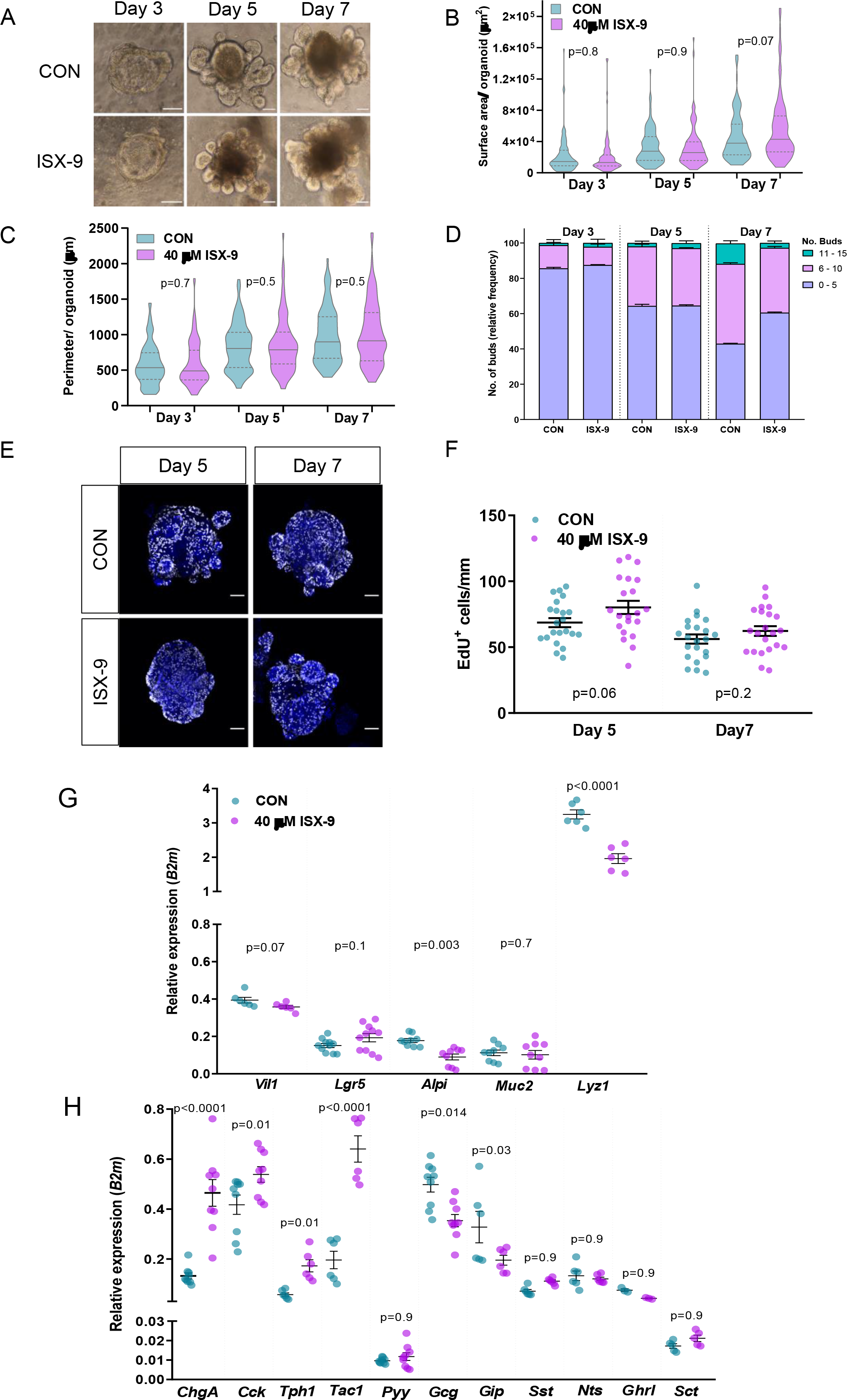
ISX-9 does not affect mouse intestinal organoid growth and specifically enriches markers of EC cells. (A) Representative brightfield images of mouse small intestinal organoids following the 48-hr ISX-9 pulse protocol. Surface area (B), perimeter (C) and number of buds (D) per control and treated organoids. (E-F) Proliferating cells in control and ISX-9 treated organoids were visualized by the incorporation of 5-ethynyl-2′-deoxyuridine (EdU) (white) and counterstain with Hoechst (blue) (E). Number of Edu^+^ cells were counted for control and ISX-9 treated organoids on day 5 (at the end of 48-hr ISX-9 treatment) and on day 7 (48 hrs after removal of ISX-9) (F). (G-H) qPCR analysis of lineage markers (G) and enteroendocrine specific markers (H) in control and 48-hr ISX-9 pulse treated organoids. Brightfield images are shown as one middle plain field of the organoid. Immunofluorescence images are shown as maximum intensity projections of a z-stack through the organoid. Scale bar: 50μm. Data are represented as mean ± SEM, except in violin plots, where data are presented as median and quartiles. Unpaired two-tailed Student’s t test (B, C, F), one way-ANOVA with Holms-Sidak multiple comparisons post hoc test (G and H). Kruskal-Wallis test with Dunn’s post hoc test (D).

To probe the effect of ISX-9 on epithelial lineages and EEC subset differentiation we analysed the expression of markers of individual cell types at the end of the differentiation protocol. In line with the growth data, expression of villin *(Vil1)*, a general epithelial marker, was unaltered between groups, as was the marker for goblet cells mucin 2 (*Muc2*) (Fig. 2G). The marker for stem cells, leucine-rich repeat-containing G-protein-coupled receptor 5 (*Lgr5*), was slightly increased but not significantly, whilst the enterocyte marker, alkaline phosphatase (*Alpi*) and the Paneth cell marker, lysozyme (*Lyz1*), were significantly reduced (Fig. 2G). Within the EEC lineage ISX-9 increased markers of I cells (*Cck*) and enterochromaffin cells (Tachykinin Precursor 1 (*Tac1*), tryptophan hydroxylase 1 (*Tph1*) and chromogranin A, (*ChgA*)) (Fig. 2H). TPH1 is the rate limiting enzyme for peripheral serotonin (5-HT) production and enterochromaffin (EC) cells are the most numerous gut endocrine cell type, producing 90% of the body’s peripheral serotonin (Gross et al., 2016). Expression of L cell markers were unchanged (*Pyy*) or reduced (*Gcg*), as was the marker for K cells (*Gip*) (Fig. 2H). The markers for D cells (*Sst*), N Cells (*Nts*), X cells (*Ghrl*) and the promiscuous marker secretin (*Sct*) were all unchanged in response to ISX-9 (Fig. 2H). Together, these data imply ISX-9 specifically increases the EEC lineage and drives cells to be fated towards enterochromaffin cells and *Cck* expressing cells rather than other mature EEC subtypes.

### ISX-9 increases *Ngn3* endocrine fated populations

To determine if we were observing specific alterations in EEC cell fate, we used lineage tracing, fluorescent activated cell sorting, single cell RNA-seq and immunostaining. First, we generated small intestinal organoids from *Tg(Ngn3-RFP)* mice which mark the *Ngn3* progenitor pool with turbo RFP (Kim et al., 2015). We saw a 5-fold increase in the percentage of *Ngn3*^+^ cells immediately after ISX-9 treatment, using flow cytometry, corroborating our expression data (Fig.3A).

To uncover the effect of ISX-9 on sub-populations of *Ngn3*^+^ cells and to attempt to identify its key regulatory mechanisms we employed single cell RNA-seq on sorted RFP^+^ cells immediately after the ISX-9 treatment. Unsupervised clustering of conserved markers between control and ISX-9 treated cells identified 9 clusters split into two major branches, endocrine and non-endocrine (Fig. 3B and C). The endocrine lineage contained 5 clusters; Early-*Ngn3* (Hi *Ngn3, Sox4*), Late-*Ngn3* (Hi *Ngn3*, *Cnot6l, Nkx2-2, Neurod1, Pax4, Runxlt1*), Endocrine progenitor (*Tac1, Chgb, Hmgn3, Cited2*), EC (*Chga, Tph1, Lmx1a, Reg4*) and EEC (*Pyy, CCK, Gcg, Nts*) (Fig. 3B, Suppl. Fig. 1A). The EC and EEC clusters are consistent with recent reports of two major branches of endocrine development, one containing EC cells and the other containing peptidergic gut hormone producing cell types (Gehart et al., 2019; Julie Piccand, 2019). The non-endocrine branch consisted of four clusters. One we designated as Goblet progenitors which exhibited strong goblet cell marker expression (eg. *Spink4*, *Muc2, Tff3*) but also contained Paneth cell markers (eg. *Lyz1*, *Defa24*). Two other clusters were identified as enterocyte progenitors on account of their expression of *Cftr and Krt19*, these clusters were distinguished from one another by one cluster expressing markers of proliferation (*Top2A, Mik67)* (Fig. 3B, Suppl. Fig. 1A). An additional hard to define cluster was designated as non-endocrine progenitors (*Fryl, Plac8, Stat1, Grk4)*.

**Figure 3:**
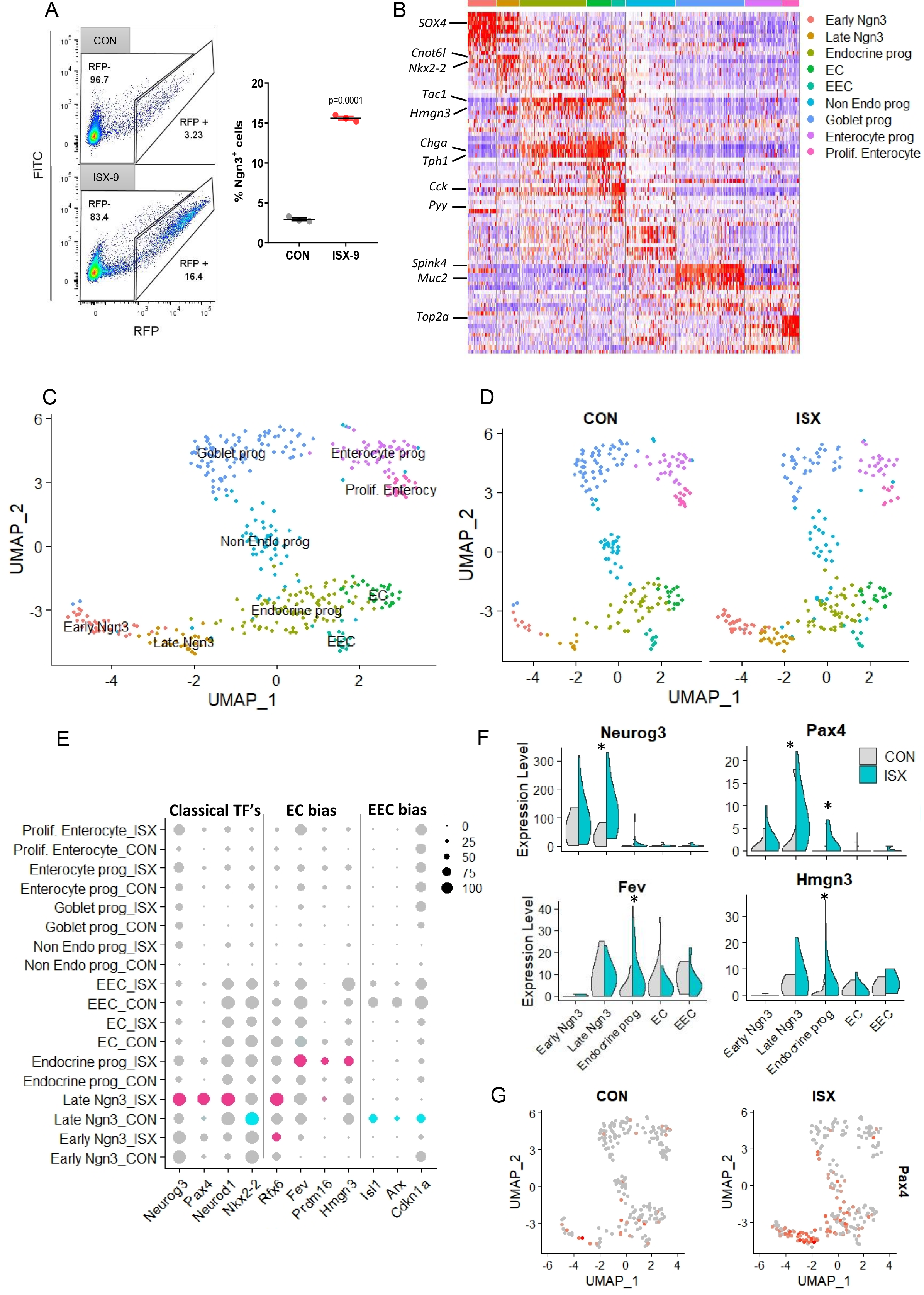
ISX-9 increases *Ngn3* endocrine fated populations with a bias toward EC cells. Intestinal organoids derived from Ngn3-RFP had a 48-hr ISX-9 treatment or left untreated. Immediately after treatment fluorescently labelled cells were sorted for further analysis. (A) Flow cytometric scatter plots of control vs. treatment and percentage of RFP^+^ cells recovered (Unpaired two-tailed Student’s t test). (B) Combined expression heatmap of top 10 conserved marker genes between control and treated cells for each cluster. (C) UMAP projection depicting clusters identified by conserved marker expression between control and treatment. (D) UMAP projection of clusters split by treatment. (E) Dot plot of transcription factors known to influence endocrine and EC cell fate comparing control versus treatment across clusters. Size of dot represents percentage of cells positive for gene within cluster, intensity of dot represents average expression of gene within cluster. (F) Violin plots of average *Neurog3, Pax4, Fev, Hmgn3* expression by cluster and treatment. (G) UMAP projections of *Pax4* expression comparing control vs. treatment. * = adj. p value for *Neurog3* (0.01), *Pax4* (late Ngn3 = 0.04, Endocrine prog = 0.0015), *Fev* (0.015), *Hmgn3* (0.03).

Examination of the cluster distributions by treatment revealed an increased proportion of cells in the developmentally early endocrine clusters, Early-*Ngn3* and Late-*Ngn3,* in response to ISX-9 (Fig. 3D). Coincidentally there was a reduction in the number of cells in the non-endocrine branch (Fig. 3D). Given the increased expression of EC markers and the I cell marker *Cck* in our original experiments, ISX-9 plausibly biases endocrine lineage cells towards these pedigrees. However, a closer examination of *Cck* expression demonstrated a low level of promiscuous expression throughout the endocrine branch in contrast to other classical gut hormones which were restricted to the EEC cluster (Fig. 3B). We explored possible mechanisms of EC bias by examining TFs known to regulate endocrine differentiation. ISX-9 increased the expression of *Ngn3 and NeuroD1* in the Late-Ngn3 cluster but notably increased both the expression and percentage of cells expressing *Pax4* in this cluster (Fig. 3E-G). Strikingly, ISX-9 increased the expression of genes associated with EC progenitor bias (*Rfx6, Fev, Prdm16 and* Hmgn3) whilst reducing genes associated with EEC bias (*Isl1, Arx and Cdkn1a*) (Fig. 3E and F, Suppl. Fig. 1B) (Gehart et al., 2019).

Together these data suggest ISX-9 increases the flux of cells through the endocrine branch in part at the expense of the non-endocrine route and increases the expression of an EC biased genetic program in these endocrine progenitors. In addition, the powerful increase in both *Pax4* expression and the percentage of *Pax4^+^* cells in the late Ngn3 cluster highlights a potentially novel mechanism of early EC bias in response to ISX-9 (Fig. 3E and G).

### ISX-9 increases the Ngn3 lineage and enriches it with functional EC cells

Next, we took advantage of a *Tg(Ngn3-cre)C1Able/J::R26-loxSTOPlox-tdRFP* (*Ngn3-Cre-RFP*) reporter transgenic line to examine the whole endocrine lineage including the *Ngn3*^+^ progenitor pool and all daughter cell types (Fig. 4A) (Luche et al., 2007; Schonhoff et al., 2004b). A 48-hr ISX-9 pulse increased the percentage of tdRFP^+^ cells analysed by flow cytometry (Fig. 4B) and doubled the number of tdRFP+ cells per organoid (Fig. 4C). Expression analysis of sorted populations confirmed our earlier findings, with *Ngn3* and *NeuroD1* being increased immediately after 48-hr ISX-9 treatment whilst *Lyz1* (Paneth) was reduced and *Muc2* (goblet) and *ChgA* (EC cell) were unaltered (Suppl. Fig. 2A). Forty-eight hours later, after removal of ISX-9, *Ngn3* was normalised, *Muc2* was unchanged, *Lyz1* remained reduced, *NeuroD1* remained elevated and as expected *ChgA* was 3-fold increased (Fig. 4D), providing evidence that ISX-9 increases the endocrine progenitor pool and biases a proportion of these cells to become specific EC cells.

**Figure 4:**
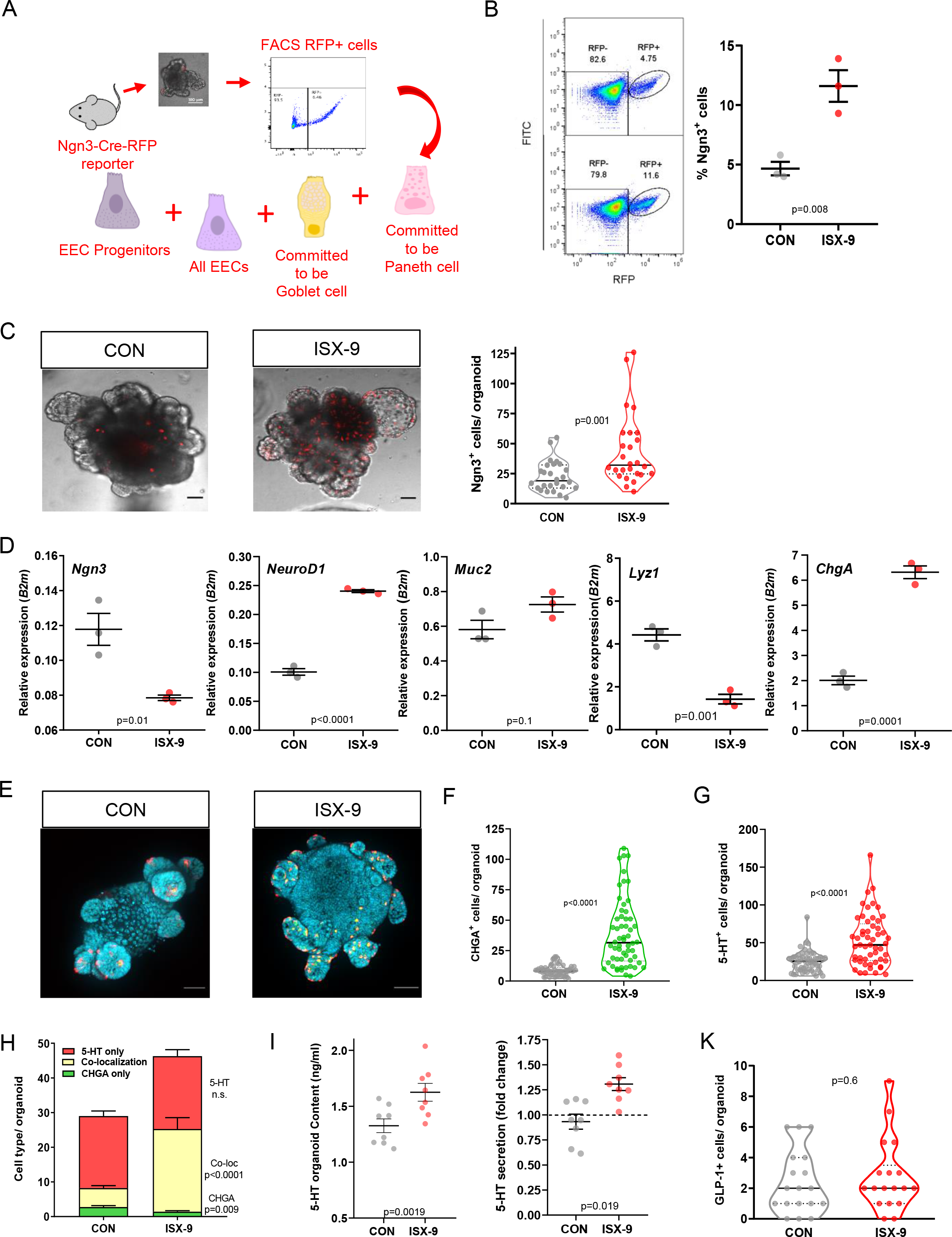
ISX-9 increases the number of Ngn3^+^ endocrine progenitors and enriches mouse intestinal organoids with and functional enterochromaffin cells. (A) Experimental design schematic. The number of RFP^+^ cells in Ngn3-Cre-RFP mouse intestinal organoids treated with a 48-hr ISX-9 pulse, was increased in comparison with controls as shown by flow cytometric analysis (B) and by counting from live-imaging (n=26) (C). (D) Expression levels of *Ngn3*, *NeuroD1*, *Muc2*, *Lyz1* and *ChgA* in RFP^+^ cells sorted from Ngn3-Cre-RFP mouse intestinal organoids. (E) Confocal images of double immunofluorescent staining of for CHGA (green) and 5-HT (red). (F-H) Quantification of singly labelled CHGA+ and 5-HT+ cells (F and G), and cells which co-localised these antigens (H) (n=53-58). (I) 5-HT content and release measured by enzyme-linked immunosorbent assay (ELISA)(n=8). (K) Quantification of GLP-1^+^ cells (n=18). Live-cell images are shown as one middle plain for brightfield and as maximum intensity projections of a z-stack for RFP. Immunofluorescence images are shown as maximum intensity projections of a z-stack. Scale bar: 50μm. All data are represented as mean ± SEM, except in violin plots where data are presented as median and quartiles. Unpaired two-tailed Student’s t test.

To further substantiate the expression data and demonstrate the programming of specific EEC subsets we generated organoids from a *CCK-iCre::R26-loxSTOPlox-eYFP* reporter mouse (*CCK*-*Cre*-*eYFP*) (D’Agostino et al., 2016; Flak et al., 2014). ISX-9 increased the number of eYFP^+^ cells per organoid by almost 3-fold (Suppl. Fig. 2B-C) and produced a similar increase in the percentage of eYFP^+^ cells by flow cytometry analysis (Suppl. Fig. 2D). In lieu of the availability of reporter mice for either chromogranin A or serotonin we used immunocytochemistry to measure the effect of ISX-9 on the EC cell lineage. Double staining of whole organoids revealed 3 cell populations: singly labelled 5-HT^+^ and CHGA^+^ cells, and cells which co-localised these antigens (Fig. 4E). ISX-9 increased the total number of 5-HT^+^ cells by 2-fold and CHGA^+^ cells by 4-fold (Fig. 4F and G). Analysis of the proportion of single versus co-localised cells revealed the increase to be driven mainly by cells expressing both 5-HT and CHGA, which are likely endocrine EC cells (Fig. 4H). ISX-9 increased organoid 5-HT content and these newly generated cells were functional, releasing measurable 5-HT in response to stimulus (Fig. 4I). ISX-9 did not alter GLP-1^+^ immunofluorescent cell number (Fig. 4K) despite *Gcg* expression being reduced, underpinning the selectivity of ISX-9’s effect.

### Enterochromaffin cell enrichment is partly dependent on ISX-9 induced calcium signalling

Mechanistically, ISX-9 promotes neuronal differentiation by stimulating intracellular calcium signalling (Schneider et al., 2008), it therefore seemed logical to consider if this was also true of ISX-9’s actions on EEC differentiation. To explore this, we used calcium fluorometry in Fura-2 AM loaded mouse organoids. ISX-9 induced a long and slow (approx. 15min) increase in mean basal to peak calcium response in 40% of cells examined (Fig. 5A and Suppl. Fig. 3A-C). All cells including those which did not respond to ISX-9, produced the expected fast spike in calcium following administration of the positive control ATP (Fig. 5A). Positionally, responders were never found at the base of the crypt buds, where Lgr5^+^ stem cells reside, but were present in various positions around and above the +4 position, and likely within the nominal transit amplifying zone (Suppl. Fig. 3D-F).

**Figure 5:**
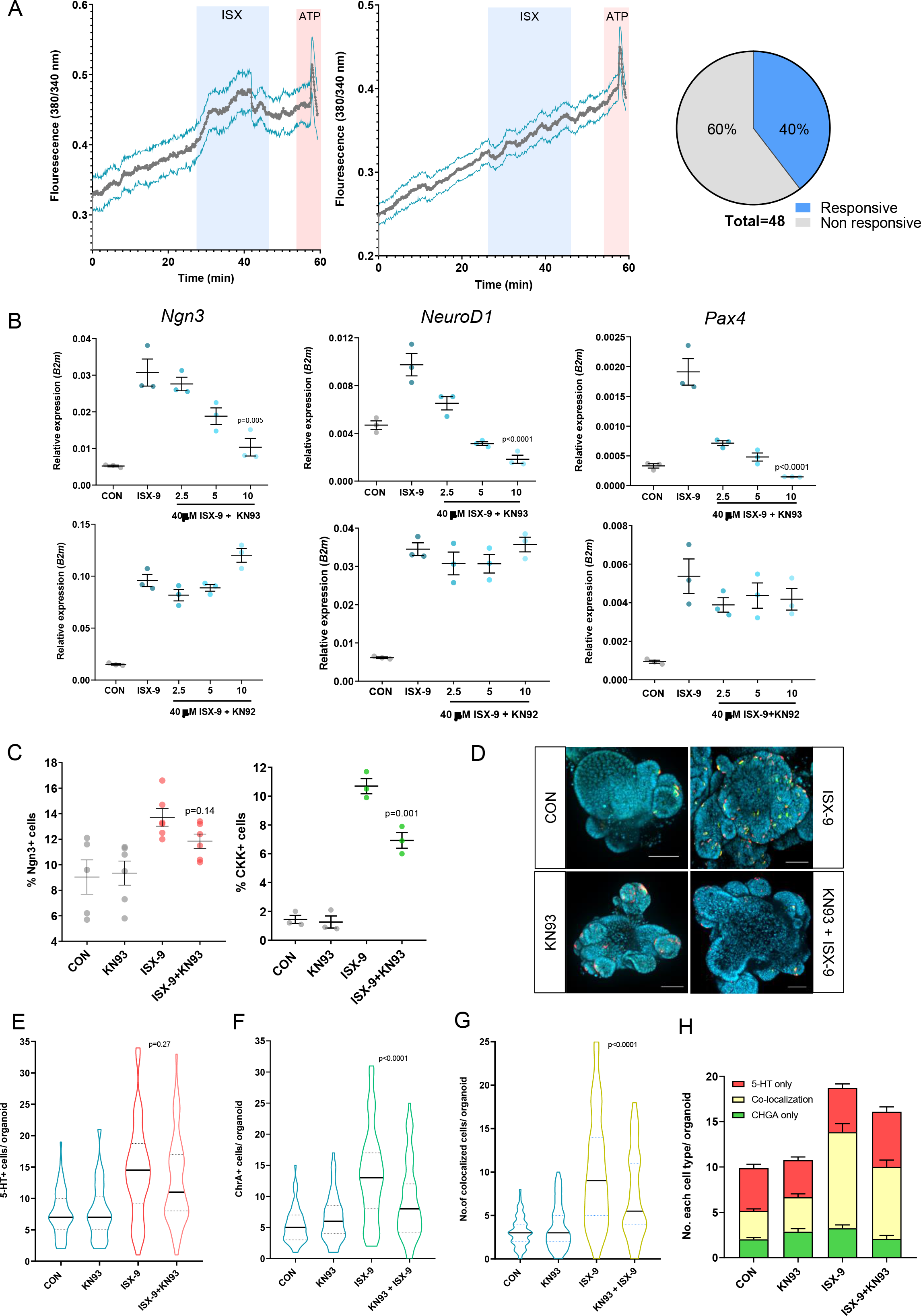
Enterochromaffin cell enrichment is partly dependent on ISX-9 induced calcium signalling. (A) Live-cell imaging of intracellular calcium in mouse small intestinal organoids using fluorescent indicator Fura-2 AM. Representative temporal plots of [Ca^2+^]i changes are shown, expressed as F340/380, in responsive cells (left) and non-responsive cells (middle) upon exposure to 40 μM ISX-9, followed by 100 μM ATP. Pie chart (right) shows the proportions of cells responsive to 40 μM ISX-9 (2 individual experiments). (B) Expression of *Ngn3*, *NeuroD1* and *Pax4* in mouse organoids after 48-hr exposure to 40 μM ISX-9 in the presence of increasing doses of KN93 (top panel) or (KN92) (bottom panel). (C) Flow cytometric analysis of Ngn3^+^ and CCK^+^ cells from Ngn3-Cre-RFP and CCK-Cre-eYFP organoids respectively, in the presence or absence of ISX-9, KN93, or combination of both, for 48 hrs. Data are representative of a single experiment with n=3. (D) Images of double immunofluorescent staining of control and ISX-9, KN93 or combination of both treated organoids for CHGA (green) and 5-HT (red). (E-G) Quantification of total 5-HT^+^ (E), total CHGA^+^ cells (F), and cells which co-localised these antigens (G) in control and ISX-9 and KN93 treated organoids. (H) Quantification of singly labelled 5-HT^+^, CHGA^+^ cells and co-localised cells (n=59-64). All confocal images are shown as maximum intensity projection of a z-stack. Scale bar: 50μm.Data are represented as mean ± SEM, except in violin plots where data are presented as median and quartiles. One-way ANOVA with Sidak’s post hoc test. (H) CHGA - CON vs ISX-9, p=0.1574; ISX-9 vs ISX-9+KN93, p=0.3217; CON vs ISX-9+KN93, p=0.9714. 5-HT - CON vs ISX-9, p=0.9878; ISX-9 vs ISX-9+KN93, p=0.1733; CON vs ISX-9+KN93, p=0.0913. Co-localisation – CON vs ISX-9, p<0.0001; ISX-9 vs ISX-9+KN93, p=0.0118; CON vs ISX-9+KN93, p<0.0001.

In neuronal progenitors ISX-9 increases calcium from both extra- and intra- cellular calcium sources (Schneider et al., 2008). We therefore chose to pharmacologically block ISX-9 induced calcium responses in organoids using the Calcium Calmodulin Kinase II enzyme inhibitor KN93. CamKII is an important intracellular calcium signalling node. Interestingly, our single cell data demonstrated the majority of *Camk2b* expressing cells were restricted to the late Ngn3 and endocrine progenitor cell populations and showed a high degree of co-localisation with *Pax4*, particularly in the Late Ngn3 progenitors (Suppl. Fig. 3G). When given in conjunction with ISX-9, KN93 dose dependently inhibited induction of *Ngn3*, *NeuroD1* and *Pax4* whilst KN93’s inactive analogue, KN92, did not (Fig. 5B). Blocking CamKII signalling reduced the effect of ISX-9 on EEC differentiation. The expansion of *Ngn3*^+^ and *Cck*^+^ cells was reduced by approximately 30-40% when KN93 was present (Fig. 5C). KN92 did not affect the expansion of *Cck*^+^ cells driven by ISX-9 (Suppl. Fig. 3H). KN93 also attenuated the increase in total CHGA^+^ cells by 50% and mildly but not significantly reduced the total number of 5-HT^+^ cells (Fig. 5D-H). As expected KN93 had little effect on the singly labelled populations of 5-HT^+^ and CHGA^+^ (Fig. 5E and F), but inhibited ISX-9 induced expansion of the co-localised population representing endocrine EC cells (Fig. 5G and H). These data reveal ISX-9 manipulates EEC differentiation in part by producing a calcium signal, likely in a population of early endocrine destined progenitors.

### Enterochromaffin cell enrichment is replicated in human terminal ileal organoids

Historically our understanding of mouse EEC differentiation has been based on knockout and lineage tracing *in vivo* studies, which were low throughput but provided key information regarding transcriptional regulation of EEC specification. The advent of organoid and single cell technologies has rapidly expanded our knowledge in this area (Gehart et al., 2019). However, there is a deficit in our understanding of human epithelial EEC differentiation. Our mouse data identified ISX-9 as a useful tool to explore features of human EEC differentiation.

We began by validating a differentiation protocol in organoids generated from terminal ileal (TI) biopsies. Organoids were stimulated to differentiate by reducing Wnt signalling for 7 days. This protocol led to reduced *LGR5* and *LYZ1* but increased *ALPI*, *VIL1* and *MUC2* (Suppl. Fig. 4). As expected, we observed strong increases in the expression of EEC markers, *NGN3*, *NEUROD1*, *CHGA*, *TPH1, GCG*, *PYY*, *SST*, *GHRL* and *NTS*, suggesting a broad augmentation of the lineage. Interestingly, *CCK* and *TAC1* were not increased during the differentiation protocol (Suppl. Fig. 4). Next, we designed an ISX-9 protocol for human TI that closely matched our mouse protocol. This consisted of a 3-day baseline period using stem cell media (WENRAS) followed by 7 days in differentiation media. A 48-hr ISX-9 pulse was delivered on day 6 (Fig. 6A). The human TI transcriptional response was remarkably similar to the mouse. *NGN3, NEUROD1* and *PAX4* expression were increased after 48-hr exposure to ISX-9 (Fig. 6B). At the end of the differentiation protocol, *ALPI*, *MUC2* and *LYZ1* expression were reduced whilst LGR5 was increased (Fig. 6C). Examination of endocrine markers following the same protocol revealed that the response to ISX-9 in human intestinal organoids mirrored the mouse. *TAC1*, *TPH1*, *CHGA* and *CCK* were increased, suggesting enrichment for EC and Cck^+^ cells, whilst markers of L cells (*PYY, GCG*), X cell (*GHRL*) and D cells (*SST*) were downregulated (Fig. 6C and D). Immunofluorescent staining confirmed EC cell enrichment, driven mainly by an expansion of cells expressing both 5-HT^+^ and CHGA^+^ (Fig. 6E-G). These new cells were functional, secreting 5-HT into the media following stimulation (Fig. 6H).

**Figure 6:**
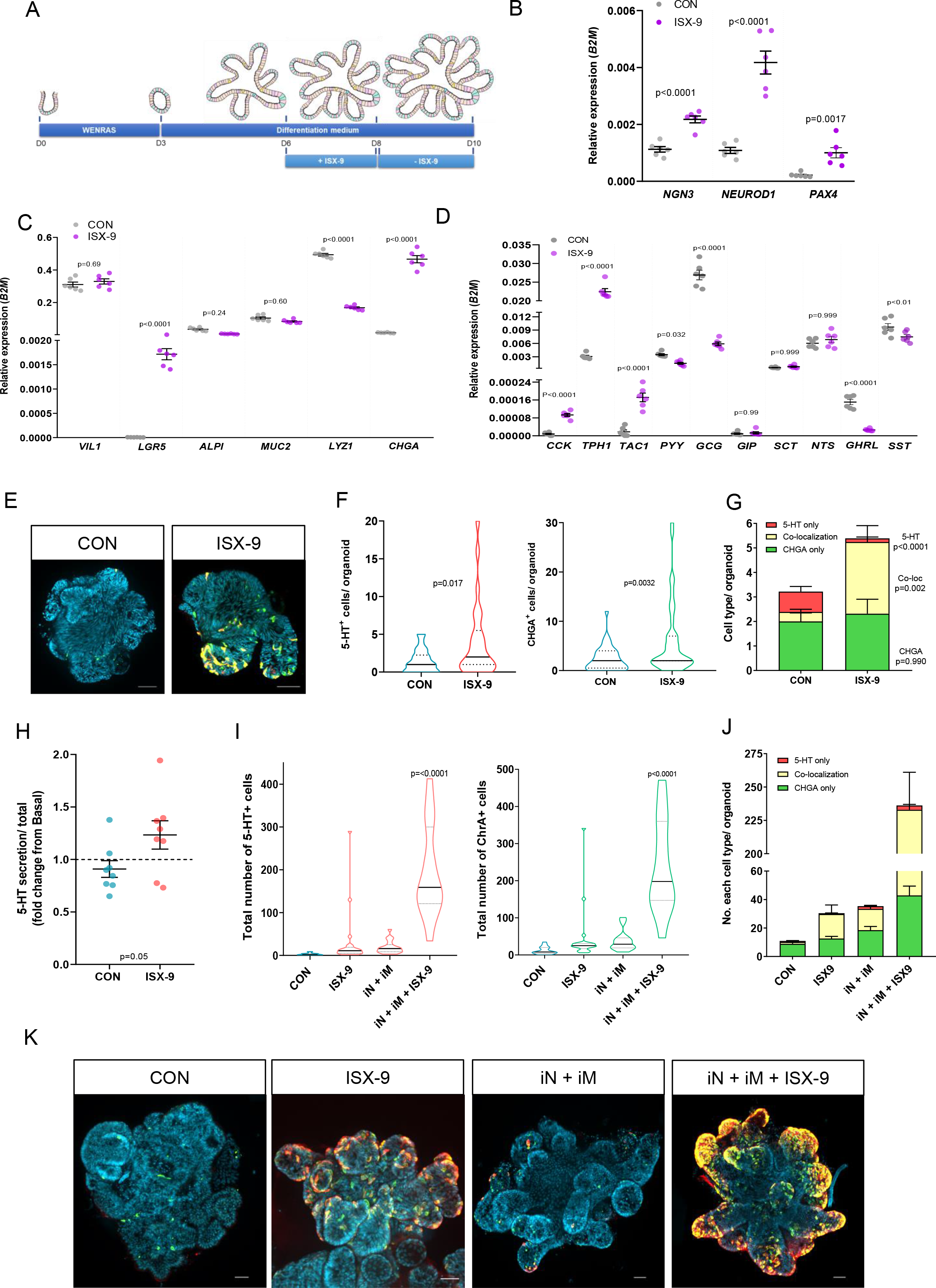
Effects of ISX-9 on human terminal ileal organoids. (A) Schematic diagram explaining the experimental paradigm of 48-hr ISX-9 pulse in human TI organoids. (B) Expression of *NGN3*, *NEUROD1* and *PAX4* in human TI organoids after 48 hrs exposure to 40 μM ISX-9. qPCR analysis of lineage markers (C) and enteroendocrine specific markers (D) in control and 48-hr ISX-9 pulse treated human TI organoids. (E) Confocal images of double immunofluorescent staining human TI organoids for CHGA (green) and 5-HT (red). (F) Quantification of total 5-HT^+^ and total CHGA^+^ cells. (G) Quantification of singly labelled 5-HT^+^ cells and CHGA^+^ cells and cells which co-localise these antigens human organoids (n=25). (H) 5-HT release from human TI organoids as measured by ELISA. (I) Quantification of total 5-HT^+^ and total CHGA^+^ cells in control human TI organoids and in organoids treated with ISX-9, a combination of iNotch and iMEK, and all three together. (J) Quantification of singly labelled 5-HT^+^ cells, CHGA^+^ cells and cells which co-localised these antigens (n=15-21). (K) Confocal images of double immunofluorescent staining of human TI organoids for CHGA (green) and 5-HT (red). All confocal images are shown as maximum intensity projections of a z-stack. Scale bar: 50μm. Data are represented as mean ± SEM, except in violin plots where data are presented as median and quartiles. Unpaired two-tailed Student’s t test (F-H). One way-ANOVA with Holms-Sidak multiple comparisons post hoc test (B-D, I and J): (J) CHGA – CON vs ISX-9, p=0.4473, ISX-9 vs iN+iM, pp=0.1062; CON vs iN+iM, p=0.0027; iN+iM vs iN+iM+ISX-9, p<0.0001. 5-HT – CON vs ISX-9, p=0.9216, ISX-9 vs iN+iM, pp=0.0006; CON vs iN+iM, p=0.0035; iN+iM vs iN+iM+ISX-9, p=0.1079. Co-localisation – CON vs ISX-9, p=0.0213, ISX-9 vs iN+iM, pp=0.9683; CON vs iN+iM, p=0.0597; iN+iM vs iN+iM+ISX-9, p<0.0001.

Overall our data point to ISX-9 programming endocrine progenitors towards an EC cell fate, prompting us to speculate whether combining ISX-9 with known stimulators of the whole EEC lineage would amplify the enrichment of EC cells. To do this, we compared the EEC transcriptional response between ISX-9; a combination of NOTCH inhibition (iNotch) and MEK inhibition (iMEK); and all three together. These inhibitors have been shown to drive EEC differentiation by inducing stem cell quiescence (Basak et al., 2017). ISX-9 and the iNotch, iMEK combination produced differential transcriptional responses. The inhibitor combination reduced *LGR5* and *LYZ1* expression and as expected increased *ALPI*, *MUC2*, *NGN3*, *NEUROD1*, *CHGA*, *TPH1*, *TAC1* and *CCK* expression (Suppl. Fig. 5). In comparison the ISX-9 response was characterised by increased *LGR5* and reduced *LYZ1* and *ALPI*. ISX-9 was equally effective as iNotch and iMEK at increasing *NEUROD1* and *CCK* but produced stronger increases in *NGN3*, *CHGA*, *TPH1* and *TAC1*, which was expected given its propensity for inducing EC cell enrichment. In combination the two protocols powerfully and synergistically enriched organoids for markers of EC cells. Immunofluorescence staining confirmed a dramatic (100-fold) enrichment for endocrine EC cells (Fig. 6I-K), providing further evidence that ISX-9 programmes early progenitors to become EC cells.

### Overexpression of *Pax4* in human ileal organoids partially mimics the effect of ISX-9 on EEC differentiation

In our original experiments *Ngn3*, *NeuroD1* and *Pax4* were identified as important transcription factors responding to ISX-9. Furthermore, at the single cell level our data highlighted a potential important role of *Pax4* in the Late-Ngn3 and endocrine progenitors which may bias cells towards an EC cell fate. To explore the role of *Pax4* we generated human TI organoids with a doxycycline (Dox) inducible *Pax4* overexpressing transgene *Tg(tetO-mPax4-IRES-mCherry)::Tg(CMV-rtTA),* using a piggyBAC system (Fig. 7A). We overexpressed mouse *Pax4*, which is 85% homologous to human *PAX4*, so that we could distinguish between endogenous and transgenic PAX4.

**Figure 7:**
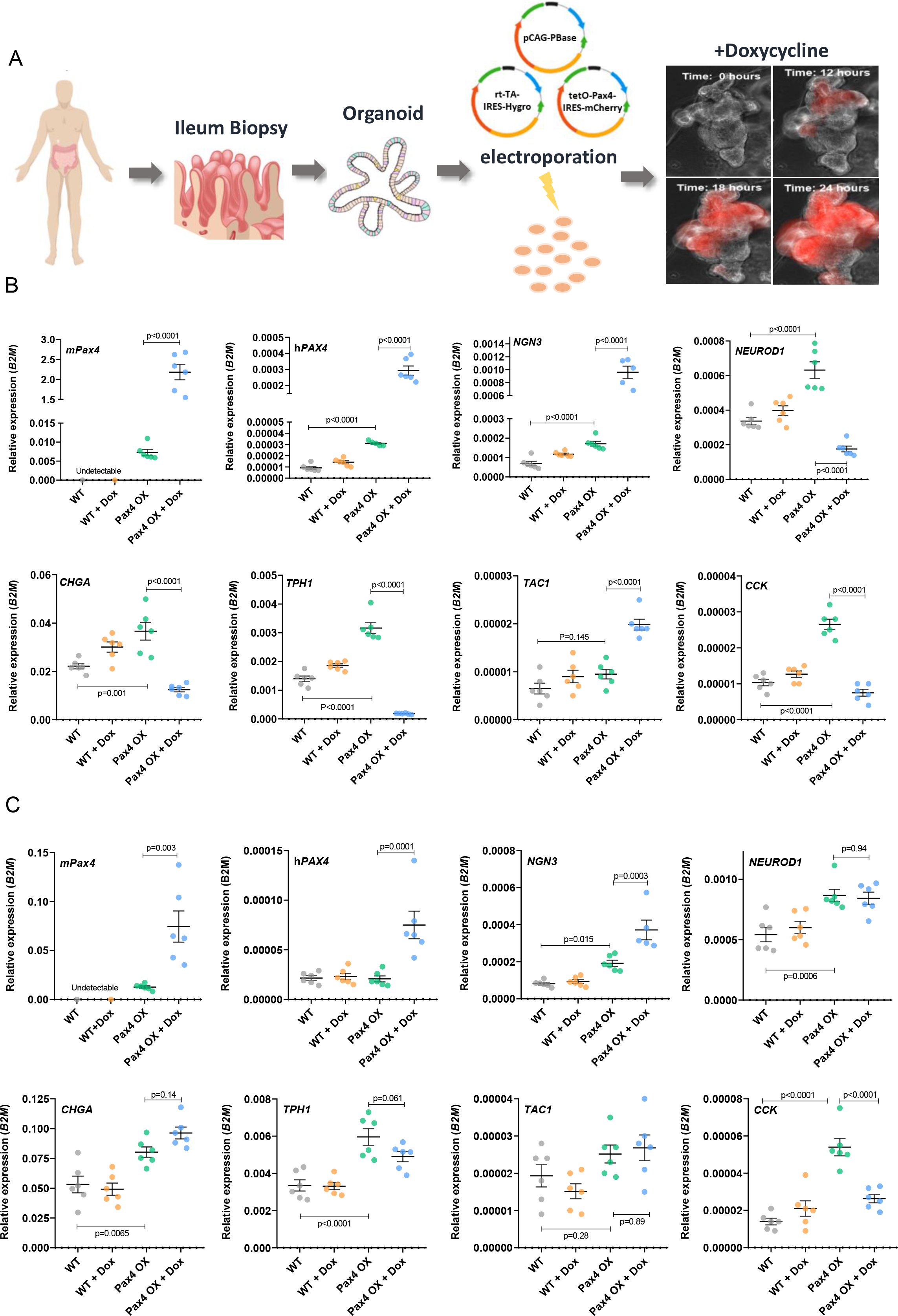
Overexpression of *Pax4* in human ileal organoids. (A) Schematic describing the production pipeline for the generation of *Pax4* overexpressing human ileal organoids. (B-C) Expression of mouse *Pax4*, human *PAX4*, *NGN3*, *NEUROD1*, EC cell (*CHGA, TPH1, TAC1*) and I cell (*CCK*) markers in wild type untreated organoids (WT), wild type organoids induced for 48 hrs with 1 μg/ml Doxycycline (WT + Dox), transgenic untreated *Pax4* human intestinal organoids (Pax4 OX) and transgenic *Pax4* human intestinal organoids induced for 48 hrs with 1 μg/ml Doxycycline (Pax4 OX + Dox) that collected for RNA extraction 48 hrs (B) or 96 hrs (C) after removal of Dox. One-way ANOVA with Tukey’s post hoc test.

Dox treatment induced RFP expression within 24 hours (Suppl. Fig. 6A, Suppl. Movie 1) correlating with a 120-fold increase in mouse *Pax4* expression, further increasing by 500-fold after 48 hours (*Pax4*^HI^) (Suppl. Fig. 6B). Doxycycline inducible transgenes have been documented to be mildly leaky (Zhu et al., 2001). Indeed, our transgenic untreated organoids exhibited a low but significant level of *Pax4* expression (3-fold) (*Pax4*^Lo^) compared to genetically identical wild-type controls maintained at the same passage. Fortuitously, the level of leaky expression was similar in magnitude to the induction by ISX-9 (Fig. 5B). This afforded us the opportunity to examine *Pax4* gene dosage on EEC differentiation. We mirrored our ISX-9 protocol by using a 3-day baseline followed by a 7-day differentiation period and gave a pulse of Dox for 48 hours on day 6. Selected transcriptional markers of epithelial cell types and EEC markers were analysed on day 10 (48 hrs after removal of Dox) (Fig. 7B). In parallel with transgenic *Pax4* expression, endogenous *PAX4* expression was also increased, 3-fold in untreated and 100-fold in Dox treated organoids, suggesting the existence of *Pax4* self-regulating pathways.

*Pax4*^Lo^ had no effect on *LGR5*, *LYZ1*, *VIL1* or *ALPI* expression but significantly increased *MUC2* expression compared to the wild type control suggesting increased goblet differentiation (Suppl. Fig. 6C). *NGN3* and *NEUROD1* expression were both significantly increased as were *CHGA* and *TPH1* (EC cells), *CCK* (I cells), *PYY* (L cells), *NTS* (N cells), *GHRL* (G cells) and *SST* (D cells) whilst *GCG* (L cells), *TAC1* (early EC cells), *GIP* (K cells) were unchanged compared to wild type (Fig. 7B and Suppl. Fig. 7A), suggesting low levels of *Pax4* overexpression induced selective specification of particular EEC subtypes including EC cells, partly mimicking the effect of ISX-9, but also inducing markers of N (*NTS*), D (*SST*) and X (*GHRL*) cells.

*Pax4*^Hi^ significantly increased *LGR5*, *LYZ1* and *VIL1* whilst reducing *ALPI* and having no effect on *MUC2* (Suppl. Fig. 6C). Unexpectedly, compared to Dox negative transgenic controls, *Pax4*^Hi^ did not enhance EEC specification, despite a 13-fold increase in *NGN3* expression (Fig. 7B). In fact, the opposite was evident; all other EEC markers (*NEUROD1*, *CHGA*, *TPH1*, *NTS*, *PYY*, *CCK*, *GHR*L, *GCG*, *SST* and *GIP*) were either strongly reduced or undetectable (Fig. 7B and Suppl. Fig. 7A), except for *TAC1* which was increased by 2-fold (Fig. 7B).

This suggested EEC differentiation was stalled by high expression of *Pax4*. To investigate this further, we measured transcriptional markers on day 12 of our protocol, 96 hrs following removal of Dox. At this time point, *Pax4*^Hi^ expression was reduced from 94-fold to a 3-fold induction (Fig. 7C). Accordingly, endogenous *PAX4* induction was halved in the *Pax4* Hi group, demonstrating *Pax4* expression was rapidly induced by Dox but was relatively slow to down-regulate following Dox removal. Elevated *NGN3* expression was reduced from 13-fold to 2-fold and *NEUROD1* was no longer suppressed (Fig. 7C). This was associated with normalisation of *TAC1* expression and a disinhibition of EC cell markers (*CHGA* and *TPH1*) (Fig. 7C). All other endocrine markers (*CCK, SST, NTS, GHRL, GCG, PYY* and *GIP*) remained supressed (Fig. 7C and Suppl. Fig. 7B). These data suggest low levels of *Pax4* expression enhance EEC specification but high levels trap EECs in an early progenitor state with an enterochromaffin cell bias. As *Pax4* expression normalises, endocrine differentiation proceeds with the appearance of EC cells. This helps explain ISX-9’s effects on EC cell enrichment and suggest upregulation of *PAX4* and *TAC1* are important.

## Discussion

Enteroendocrine cells respond to diverse signals in the luminal environment including nutrients, microbial metabolites and pathogens. They play a central role in integrating these complex signals and altering physiology by modulating epithelial, immune, neuronal and hormonal functions. These features make the enteroendocrine system a potential target for treating multiple conditions, notably attention has focused on metabolic diseases (Sam et al., 2012). An increasingly important feature of EECs is their plasticity, which hypothetically could be appropriated to alter their density and/or functional characteristics for therapeutic gain (Tsakmaki A, 2017). This is exemplified by two studies, which increased the secretory lineage *in vivo* using either a Notch or Rho-associated coiled-coil-containing protein kinase (ROCK) inhibitor (Petersen et al., 2018; Petersen et al., 2015). The increased EEC density included GLP-1-producing L cells, promoting glucose control and reducing hyperglycaemia in models of diabetes. The caveat to these proof of principle studies is their broad effect across the secretory lineage. Targeting specific EEC differentiation pathways might be a more suitable approach but requires a deeper understanding of the regulatory networks controlling EEC specification, particularly in the human epithelium, which had been difficult to study before the advent of organoid technology.

We used ISX-9, a small molecule activator of *NeuroD1*, to further investigate EEC differentiation in mouse and human intestinal organoids. In the gut, *NeuroD1* is downstream of *Ngn3* in endocrine progenitors and its deletion reduces the number of CCK and secretin cells in mice (Schonhoff et al., 2004a), potentially offering the opportunity to modulate specific EEC cell fates using ISX-9. Indeed, in our initial experiments ISX-9 strongly increased classical TFs known to be key members of the regulator network controlling EEC differentiation (*Ngn3*, *NeuroD1* and *Pax4*), but did not affect *Arx*, suggesting specificity. Analysis of organoids derived from transgenic reporter mice or immunostained for serotonin, revealed ISX-9 increased markers of endocrine progenitor development and demonstrated an enrichment of functional terminally differentiated EC cells. At single cell resolution we were able to corroborate the presence of two major developmental endocrine trajectories, peptidergic (gut hormone) versus enterochromaffin and identify developmentally earlier endocrine clusters. Analysis of ISX-9 treated cells provided evidence of increased cells in the endocrine branch at the expense of the non-endocrine goblet/enterocyte lineages. A notable increase of Pax4 expression in the Late-Ngn3 population coupled with an activation of a genetic program biased towards EC differentiation could explain the enrichment in functional EC cells. Importantly, this appeared to be a genuine manipulation of lineage fate and not an alteration in organoid growth. Equally, organoid proliferation and *Lgr5* expression were unaltered, in contrast to a previously published paradigm which promotes EEC lineage differentiation by inducing Lgr5 stem cell quiescence using NOTCH and MEK inhibitors (Basak et al., 2017).

The triggering of neuronal differentiation by ISX-9 is calcium dependent (Schneider et al., 2008). Similarly, its effect on the endocrine lineage was partly dependent on calcium signalling. Inhibition of the intracellular calcium signalling node, CamKII, attenuated EC and *Cck^+^* cell enrichment and blocked induction of *Ngn3*, *NeuroD1* and *Pax4* expression. This chimes with recent data in Drosophila, showing activation of the mechanosensitive receptor Piezo in a population of mid-gut endocrine progenitor cells increases EEC differentiation through cytosolic Ca^2+^ (He et al., 2018). In mice, Piezo2 is found in a sub-population of EC cells which release serotonin in response to stretch (Alcaino et al., 2018). Future work will be aimed at identifying in which cells ISX-9 stimulates calcium and whether calcium signalling is an important regulator of homeostatic endocrine differentiation in the gut.

Most data relating to EEC differentiation has been gathered in mice or using mouse tissues. Comparatively little is known about the networks controlling endocrine cell fate in the human gut epithelium, but they are assumed to be conserved to some degree, stemming largely from the study of patients with mutations in *NGN3* who exhibit intractable malabsorptive diarrhoea due to loss of EECs and a handful of recent studies in organoids (German-Diaz et al., 2017; Pinney et al., 2011; Rubio-Cabezas et al., 2011). In these *in vitro* studies, a pulse of transgenic *NGN3* in iPSCs-derived human organoids mostly recapitulated the predicted endocrine repertoire (Sinagoga et al., 2018). The same transgenic construct has been deployed in organoids derived from tissue resident stem cells and similarly promoted EEC differentiation (Chang-Graham et al., 2019). Our data shed further light on human EEC differentiation and its similarity to mouse. Notably, ISX-9 produced identical effects on the EEC lineage between species. *NGN3*, *NEUROD1* and *PAX4* up-regulation was associated with selective increases in *CCK*, *CHGA*, *TPH1* and *TAC1*. Whole organoid immunostaining confirmed enrichment for EC cells double positive for 5-HT and CHGA and these cells functionally released 5-HT. Our data have several implications; they highlight how similar EEC specification is between mouse and human and that ISX-9 alone or in combination with iNotch and iMEK can be used to enrich human organoids with EC cells allowing functional exploration of these rare and difficult to study cells. EC cells modulate GI motility, bone formation, hepatic gluconeogenesis, thermogenesis, insulin resistance, and regulation of fat mass (Yabut et al., 2019). Understanding how these cells function in the human has important implications.

Our data highlighted the potential importance of *Pax4*, its induction in the late Ngn3 population seemed to offer a developmentally early mechanistic explanation for EC cell enrichment by ISX-9. However, low level constitutive expression produced a generalised increase in EEC lineage markers, notable exceptions were *GCG* and *GIP*. We also observed a surprising and powerful upregulation of *NGN3*, suggesting reciprocal regulation between *PAX4* and *NGN3*. The upregulation of *MUC2* and *LYZ1* might also be explained by this *NGN3* upregulation, as 15% of goblet and 40% of Paneth cells are derived from Ngn3 progenitors (Schonhoff et al., 2004b). Strikingly, induction of high levels of *Pax4* expression completely inhibited endocrine differentiation, trapping EEC differentiation at an early stage. Unexpectedly, induced transgenic *Pax4* was persistent remaining upregulated by 3-fold 96 hours after doxycycline removal, although this was reduced from a peak induction of 94-fold. Interestingly, at this time point all markers of peptidergic EECs (*CCK, SST, NTS*, *GHRL*, GCG and PYY) remained supressed, but markers of mature EC cells (*CHGA and TPH1*) were now disinhibited, and *TAC1*, an early EC cell marker, was no longer upregulated. We drew several conclusions from this. Firstly, our data are consistent with the existence of two major lineages of EECs in the human, as recently described in the mouse and corroborated by our single cell data (Gehart et al., 2019; Julie Piccand, 2019). One lineage giving rise to an early appearing EC cell population and the other to peptidergic producing cells, which generally appear later than EC cells. Secondly, *PAX4*, as in the mouse, is upstream of *NEUROD1* and likely marks an early endocrine progenitor cell. Thirdly, understanding the network of TFs controlling EEC differentiation is complicated by their promiscuity and the need to appreciate the timing and level of expression. Lastly, it seems plausible that ISX-9 enriches the ECC lineage in part by its effects on *PAX4* expression which may represent an increase in the endocrine pool biased towards an EC cell fate.

Finally, our data add to a handful of studies suggesting small molecules could be found to selectively control EEC cell fate/specification with a view to treat various clinical conditions. However, our data also highlight the difficulties this approach faces. Understanding to what degree the targeted lineage creates a deficit in another and whether this induces unwanted physiological effects, is of key importance. This seems a likely event when intervening downstream of *NGN3*, where one EEC type may be enriched at the expense of another. This could be mitigated by combining selective EEC targeting with a more generalised EEC lineage activator. Realising this potential will require a much deeper understanding of human EEC lineage specification aided by the organoid platform.

## Supporting information

All supplemental figures

## Acknowledgements

For providing us with intestinal tissues from transgenic animals for the generation of small intestinal organoids we would like to thank Prof Anne Grappin-Botton (Neurog3-RFP mice), Dr Mathieu Latreille (*Tg(Neurog3-cre)C1Able/J::R26-loxSTOPlox-tdRFP* mice) and Dr Giuseppe D'Agostino (*CCK-iCre::R26-loxSTOPlox-eYFP* mice). We would also like to thank Dr Calvin Kuo and Dr Hans Clever for providing us with R-Sponding1-producing cell line and L-Wnt3A cells, respectively, and Dr Bon-Kyoung Koo for supplying the piggyBAC system for the generation of doxycycline induced Pax4/RFP overexpressing human ileal organoids. Finally, we would like to thank the staff in the Nikon Imaging Centre and in BRC Flow cytometry core at King’s College London for all their help.

## Author contributions

A. T. and P. F. P. helped in the design of experiments, collected data and contributed to the writing of the manuscript. P.P and B.H.H provided human biopsies for the isolation of crypts. G. A. B. wrote the manuscript and managed the project.

## Figure Legends

**Supplementary Figure 1: Extended single cell analysis, Related to Figure 3.** (A) Violin plots of archetypal gene expression for the 9 clusters identified in Fig. 3C. (B) Average gene expression for transcription factors involved in EEC differentiation (*NeuroD1, Nkx2.2, Rfx6, Prdm16, Isl1, Arx*) focusing on endocrine clusters and split by treatment (CON, 40 μM ISX-9).

**Supplementary Figure 2: ISX-9 increases the number of Ngn3^+^ endocrine progenitors and enriches mouse intestinal organoids with CCK-expressing cells, Related to Figure 4.** (A) Expression levels of *Ngn3*, *NeuroD1*, *Muc2*, *Lyz1* and *ChgA* in RFP^+^ cells sorted from Ngn3-Cre-RFP mouse intestinal organoids treated with or without ISX-9 for 48 hrs. 48-hr ISX-9 pulse increased the number of eYFP^+^ cells in CCK-Cre-eYFP intestinal organoids as shown by live-cell imaging (B-C) and by flow cytometry (D). (B) Live-cell fluorescence microscopy images of control and ISX-9 treated CCK-Cre-eYFP intestinal organoids. eYFP labels CCK^+^ cells (I cell) (n=24). (C) quantification of (B). Live-cell images are shown as one middle plain for brightfield and as maximum intensity projections of a z-stack for eYFP. Scale bar: 50μm. All data are represented as mean ± SEM, except in violin plots where data are presented as median and quartiles. Unpaired two-tailed Student’s t test.

**Supplementary Figure 3: Enterochromaffin cell enrichment is partly dependent on ISX-9 induced calcium signalling, Related to Figure 5.** (A) Live-cell imaging of intracellular calcium in mouse small intestinal organoids using fluorescent indicator Fura-2 AM. Representative temporal plots of [Ca2+]i changes (expressed as F340/380) in 8 regions of interest corresponding to responsive cells (A) and 23 regions of interest corresponding to non-responsive cells (B) upon exposure to 40 μM ISX-9, followed by 100 μM ATP. (C) Mean peak amplitude of cells responsive and non-responsive to 40 μM ISX-9 (Unpaired two-tailed Student’s t test). (D) Image of Fura-2 AM ratio (F340/380) shown in pseudocolour, before start of experiment. (E) Regions of interest selected on bud 1. (F) Regions of interested selected on bud 2. (G) UMAP projections showing Pax4-(blue), CamK2b-(red) expressing cells and co-localised cells (purple) preferentially expressed in the late Ngn3 and endocrine progenitor clusters. (H) Flow cytometric analysis of CCK^+^ cells from CCK-Cre-eYFP organoids treated with ISX-9, KN92 or combination of both 48 hrs. Data are representative of a single experiment with n=3. All flow plots show the gating and present both positive and negative sorted populations (One-way ANOVA with Sidak’s post hoc test).

**Supplementary Figure 4: Characterization of human TI organoids**. Expression of lineage markers (*LGR5*-marker for stem cells, *LYZ1*-marker for Paneth cells, *VIL1*-a general epithelial marker, *ALPI-* marker of enterocytes, *MUC2* -marker for goblet cells), enteroendocrine specific markers (*CCK*-marker for I cells, (*CHGA*,*TPH1*, *TAC1*)-markers for EC cells, (*PYY*, *GCG*)-markers of L cells, *GIP*-marker of K cells, *GHRL*-marker of X cells, *NTS*-marker of N cells, *SST*-marker of D cells) and the TFs, *NGN3* and *NEUROD1,* in human TI organoids cultured in stem cell and differentiation medium. Unpaired two-tailed Student’s t test.

**Supplementary Figure 5: Transcriptional responses of ISX-9 combined with iNotch and iMEK in human terminal ileal organoids, Related to Figure 6.** qPCR analysis of lineage markers (*VIL1*, *LGR5*, *LYZ1*, *ALPI*, *MUC2*), TFs (*NGN3*, *NEUROD1*) and EC cell (*CHGA*, *TPH1*, *TAC1*) and I cell (*CCK*) markers in human TI organoids treated with 40 μM ISX-9, a combination of 500 nM PD0325901 (inhibition of MEK signalling-iMEK) and 10 μM DAPT (inhibition of Notch signalling-iNotch), or a combination of all three (ISX9+ iNotch iMEK) for a 48-hr pulse. One-way ANOVA with Sidak’s post hoc test.

**Supplementary Figure 6: Further characterization of human ileal organoids overexpressing***Pax4***, related to Figure 7.** (A) Time-lapse video showing the induction of RFP expression in transgenic Pax4 human intestinal organoids with 1 μg/ml Doxycycline within 24 hours (left) which correlated with a 120-fold increase in mouse *Pax4* expression (right). (B) Quantification of mouse *Pax4* expression after treatment with 1 μg/ml Doxycycline for 48h (C) qPCR analysis of lineage markers (LGR5, LYZ1, VIL1, ALPI, MUC2) in wild type untreated organoids (WT), wild type organoids induced for 48 hrs with 1 μg/ml Doxycycline (WT + Dox), transgenic untreated Pax4 human intestinal organoids (Pax4 OX) and transgenic Pax4 human intestinal organoids induced for 48 hrs with 1 μg/ml Doxycycline (Pax4 OX + Dox). Samples were collected for RNA extraction 48 hrs after removal of Dox. One-way ANOVA with Sidak’s post hoc test.

**Supplementary Figure 7: Further characterization of human ileal organoids overexpressing *Pax4*, Related to Figure 7.** Expression of EEC markers *SST, NTS, GHRL, GCG, PYY*, and *GIP* in wild type untreated organoids (WT), wild type organoids induced for 48 hrs with 1 μg/ml Doxycycline (WT + Dox), transgenic untreated Pax4 human intestinal organoids (Pax4 OX) and transgenic Pax4 human intestinal organoids induced for 48 hrs with 1 μg/ml Doxycycline (Pax4 OX + Dox), collected for RNA extraction 48 hrs (A) or 96 hrs (B) after removal of Dox. One-way ANOVA with Sidak’s post hoc test.

---

**Key resources table:**
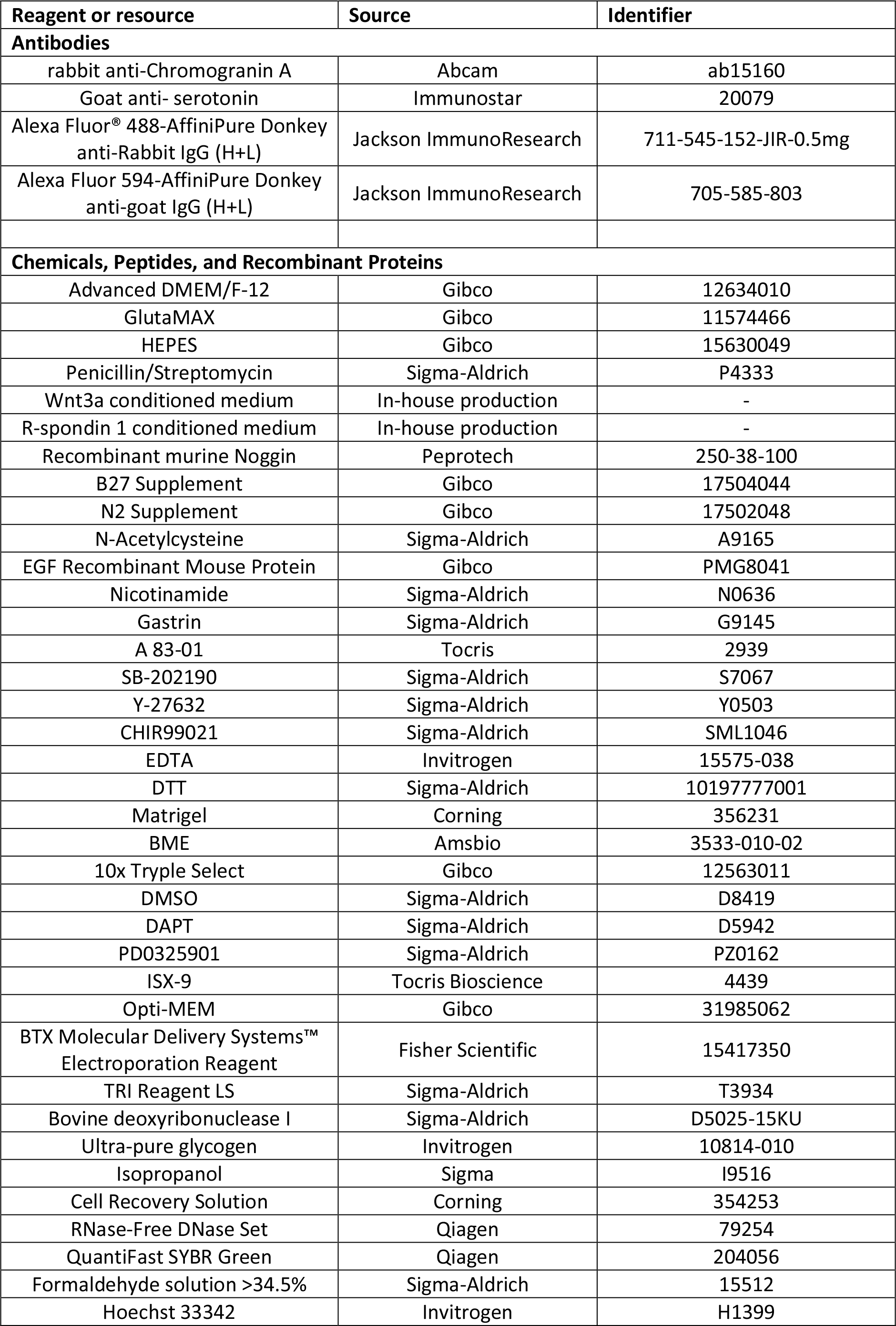
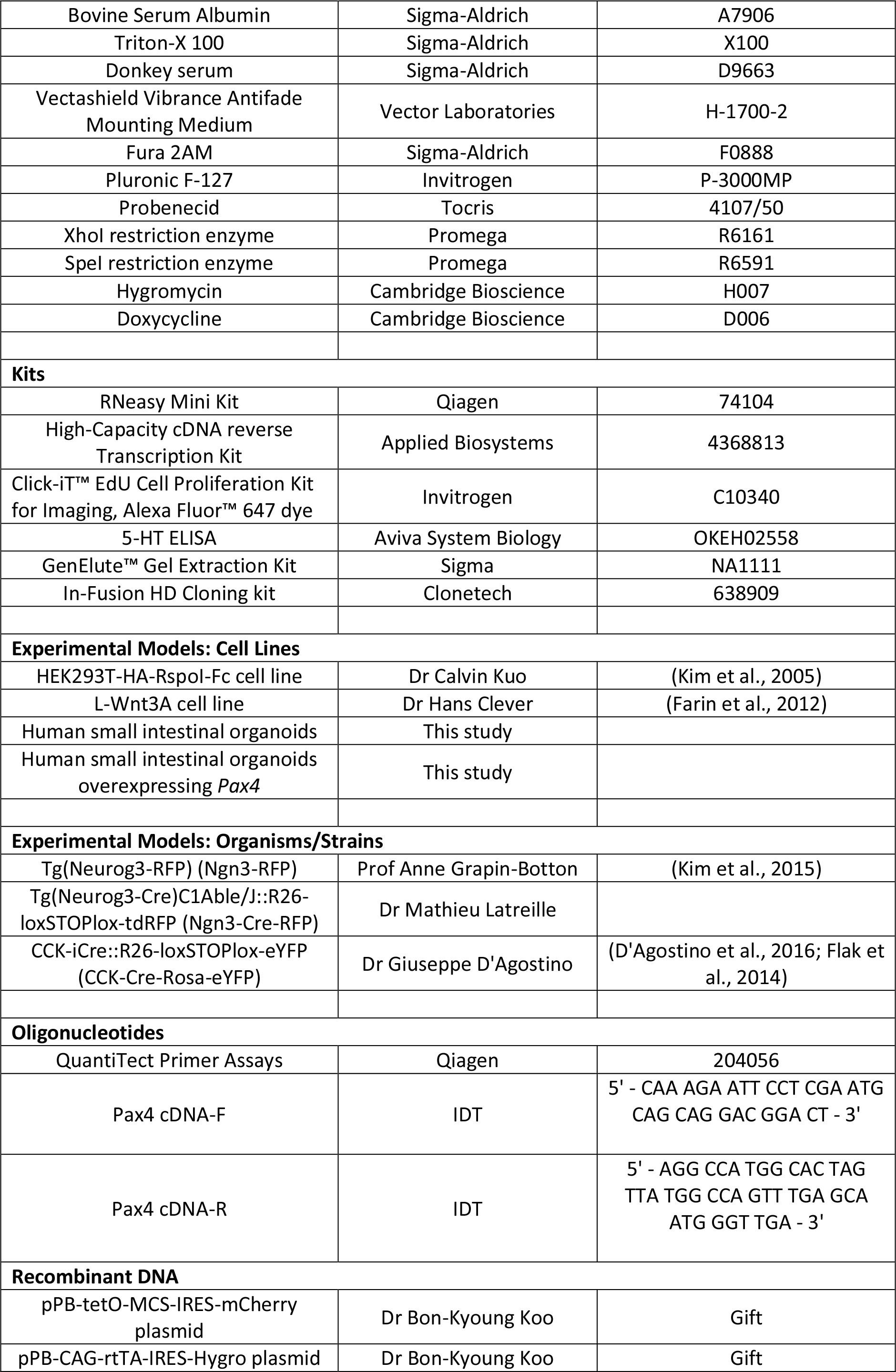
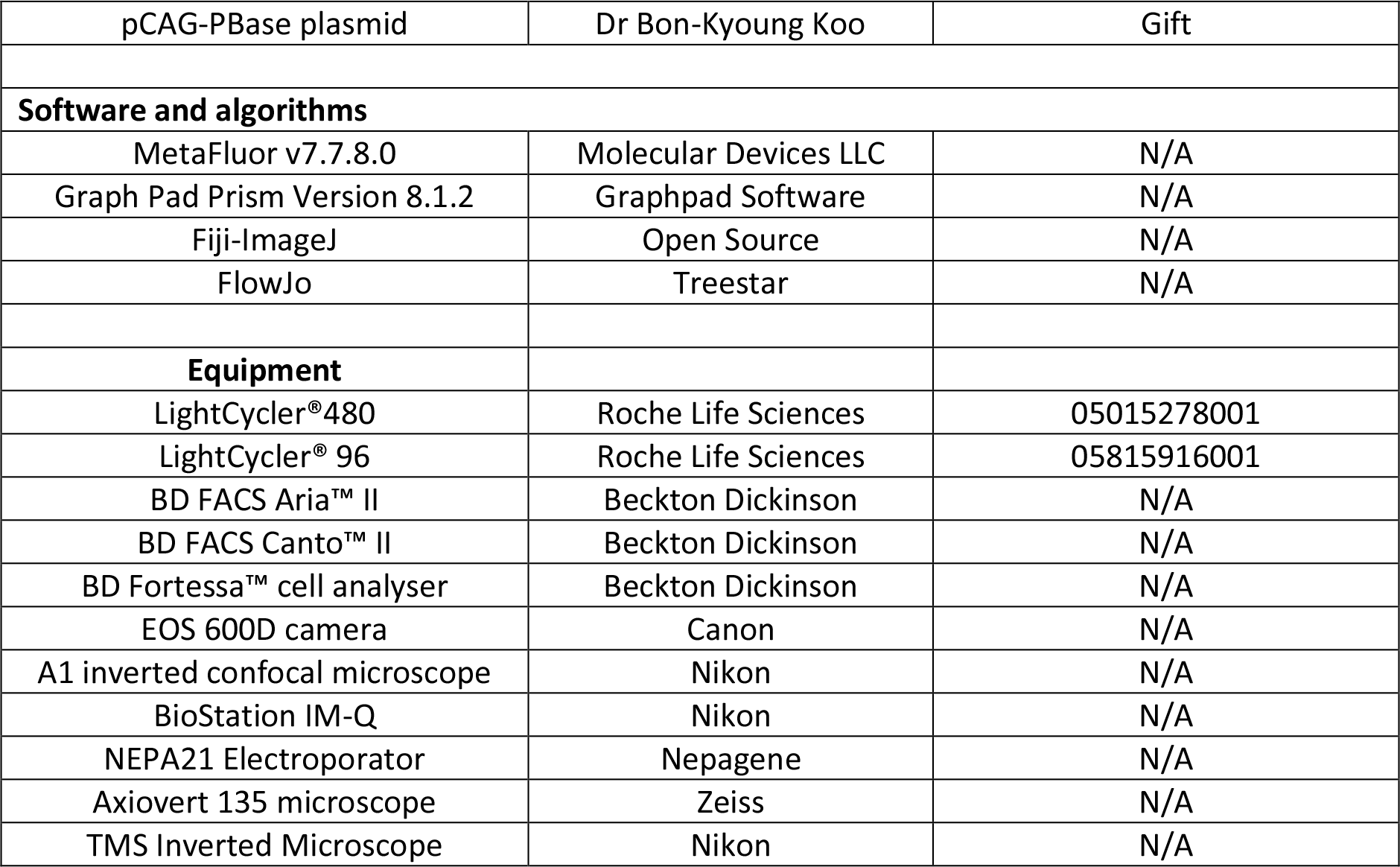

## Crypt isolation and mouse intestinal organoid culture

Mouse small intestines were harvested and cleaned with cold PBS and separated into 2 parts: duodenum (proximal 5cm), jejunum and ileum. Each part was cut longitudinally, and villi were scrapped with a glass slide. The tissue was cut with scissors into 2×2 mm pieces and repeatedly washed. Subsequently, the tissue pieces were incubated with 2 mM EDTA in PBS for 45min, in a rotator at 4 °C. After removal of EDTA, vigorous shaking in cold PBS lead to release of crypts. Crypts were further washed in PBS, passed through a 40 μm cell strainer, pelleted and resuspended in BME. Crypts were plated in 48-well plates, with 200 crypts per 25 μl of BME. The BME was polymerized for 15 min at 37°C and stem cell growth medium (WENR) supplemented with 10 μM Y-27632 was overlaid. WENR medium consists of Advanced DMEM/F12, 2 mM GlutaMAX, 10mM HEPES, 100 units/mL penicillin/streptomycin, 50 ng/mL EGF, 1× B27, 1× N2 supplements (all from Gibco), 1.25 mM N-Acetylcysteine (Sigma-Aldrich), 100 ng/ml Noggin (Peprotech), 50% Wnt3A conditioned medium and 10% R-spondin-1 conditioned medium (both in house production). Three days later, medium was changed into differentiation media (ENR) with no Wnt3A or Y-27632. Organoids were passaged once a week by mechanical dissociation, at a 1:3 split ratio. Plated organoids were maintained in a CO2 incubator with 5% CO2 and the media were changed every other day.

Treatment of mouse small intestine organoids with ISX-9 started 3 days after passaging. For dose-response experiments organoids were treated with 2 μM, 20 μM, 40 μM and 80 μM ISX-9 (Tocris Bioscience) for 48 hrs. For time-course experiments, organoids were treated with 40 μM ISX-9 for 24 hrs, 48 hrs, and 96 hrs and samples were collected for RNA extraction at the end of each timepoint. Treatment with 10 μM KN93 or 10 μM KN92 (an inactive analogue of KN93) in the presence or absence of ISX-9 was also performed for 48 hrs. For the rest of the experiment’s organoids were treated with 40 μM ISX-9 for 48 hrs and then ISX-9 was removed for another 48 hrs (48-hr ISX-9 pulse).

## Generation and culture of human terminal ileal organoids

Human terminal ileum crypts were isolated from biopsies acquired from patients undergoing colonoscopy at Guy’s and St Thomas’ NHS Foundation Trust with their informed consent. Biopsies were washed in cold PBS until the supernatant was clear. Following 10 minutes incubation at room temperature with 10 mM DTT, the biopsies were incubated with 8 mM EDTA in PBS, and placed in a rotator for 1 hr at 4 °C. At the end on the incubation, EDTA was removed and crypts were released with vigorous shaking in cold PBS. Crypts were further washed in PBS, pelleted and resuspended in Matrigel, in the same density as mouse crypts. Human intestinal crypts embedded in Matrigel were overlaid with stem cell growth medium (WENRAS) supplemented with 10 μM Y-27632 and 5 μM CHIR99021. Human stem cell growth medium in comparison with the mouse one described above, additionally contained 10 nM gastrin (Sigma-Aldrich), 500 nM A83-01 (Bio-techne), 10 μM SB202190 (Sigma-Aldrich) and 10 mM Nicotinamide (Sigma-Aldrich). 3 days after isolation or splitting, Y-27632 and CHIR99221 were removed from the medium and organoids were either maintained in WENRAS or transferred into differentiation medium for setting up experiments. For differentiation of human ileal organoids Wnt3A conditioned medium was reduced from 50% to 15% and SB202190 and nicotinamide were withdrawn from the medium. Differentiation medium was used for 7 days, keeping the organoids in culture for 10 days. 5 days after passaging, human terminal ileal organoids were treated with 40 μM ISX-9 for 48 hrs and then ISX-9 was removed for another 48 hrs (48-hr ISX-9 pulse), except if it is stated differently. MEK signalling was inhibited with 500 nM PD0325901 (Sigma-Aldrich), and Notch signalling with 10 μM DAPT (Sigma-Aldrich). Combination of both inhibitors with ISX-9 was given in a 48-hr pulse. All control organoids were treated with vehicle.

## RNA extraction and real time quantitative PCR

Total RNA was isolated from organoids (released from Matrigel or BME with Cell Recovery solution (Corning)) using RNeasy Mini Kit (Qiagen) according to manufacturer’s instructions. On-column DNase digestion was performed for removing any residual genomic DNA (Qiagen). RNA extraction from sorted cells was performed using TRI Reagent LS according to manufacturer’s instruction. cDNA was generated using the High-Capacity cDNA reverse Transcription Kit (Applied Biosystems). Real time qPCR was performed using QuantiTect primers and QuantiFast SybrGreen PCR kit (both from Qiagen) on a LightCycler 480 or LightCycler 96 (Roche). Relative gene expression levels were calculated by averaging the Ct values of technical duplicates for each biological sample and normalizing to the expression of the housekeeping gene Beta-2-Microglobulin.

## Evaluation of mouse small intestinal organoid growth

Growth and convolutedness of organoids was evaluated by collecting brightfield images of control and ISX-9 treated organoids with EOS 600D Nikon camera on an TMS inverted microscope (Nikon) on day 3, 5 and 7 in culture. Surface area, perimeter and number of buds were measured using ImageJ software.

## Immunofluorescent staining of organoids

Mouse or human small intestine organoids were fixed for 45 min in 4% formalin and then washed with 2% BSA in PBS. Subsequently, organoids were blocked with blocking buffer consisting of 2% BSA and 5% donkey serum, and permeabilized with 0.5% Triton-X for one hour at room temperature. Primary antibody incubation was done in a rotor overnight at 4°C with primary antibodies diluted in blocking buffer. Primary antibody used were rabbit polyclonal anti-Chromogranin A (1:800: Abcam) and goat polyclonal anti-serotonin (1:100; Immunostar). The next day, organoids were washed and incubated with secondary antibodies; Alexa Fluor 488 Donkey Anti-Rabbit or Alexa Fluor 594-Donkey anti-goat (1:500; Jackson ImmunoResearch) for 1 hour at room temperature. Nuclear counterstaining was performed in parallel with the secondary antibody incubation using Hoechst 33342 (1:2000; Invitrogen). After washing, organoids were mounted with Vectashield Vibrance Antifade Mounting Medium (Vector Laboratories).

The fraction of proliferating cells in control and ISX-9 treated mouse intestinal organoids was determined using the Click-iT Edu Cell proliferation kit, Alexa Fluor™ 647 dye (Invitrogen). Organoids were pre-incubated for 1 hr with 10 μM EdU, then fixed and permeabilized, and Edu positive nuclei were labelled according to manufacturer’s instructions.

## Immunofluorescent imaging

Live-cell fluorescence imaging was carried out in control and ISX-9 treated Ngn3-RFP, Ngn3-Cre-RFP, and CCK-Cre-Rosa-eYFP organoids. A continuous z dimension stack of RFP or eYFP fluorescence and brightfield images was obtained, while organoids were still embedded on BME, using an A1 inverted confocal microscope (Nikon). Images of whole-mount organoids stained for chromograninA and serotonin, were also capture with an A1 inverted confocal microscope (Nikon). Image analysis was performed using either Nikon Elements or Image J software. Time-lapse fluorescent microscopy of induced with Dox human intestinal organoids for overexpression of *Pax4*, was performed with a BioStation IM-Q (Nikon).

## Flow cytometry analysis and fluorescent activated cell sorting (FACS)

Control and ISX-9 treated Ngn3-RFP, Ngn3-Cre-RFP and CCK-Cre-Rosa-eYFP organoids were dissociated into single cells with mechanical disruption after 5 min incubation with TryplE Express (Gibco) at 37°C. After washes with PBS the cells were passed through a 40 μm cell strainer and resuspended in Advanced DMEM/F12 medium with 4 μg/mL DNase, 10 μM Y-27632 and 2 mM EDTA. 1μg/ ml DAPI was added to the cell suspension to label dead cells. Viable cells were analysed in a BD FACS Canto™ II (Beckton Dickinson). For RNA extraction of Ngn3^+^ cells from control and ISX-9 treated Ngn3-Cre-RFP organoids, organoids were first dissociated into single cells as described above and immediately sorted using a BD FACS Aria™ II (Beckton Dickinson).

## Sample preparation for scRNA-seq

Ngn3-Cre-RFP organoids were treated with ISX-9 for 48 hrs, collected and processed for fluorescent activated cell sorting as described above. DAPI was added just before sorting. RFP-positive, DAPI-negative cells were sorted into 384-well plates containing 384 unique molecular identifier (UMI) barcode primer-sets using a FACS Aria™ II (Beckton Dickinson). Samples in plates were centrifuged and stored at −80° C. Samples were then processed by Single Cell Discoveries B.V. according the SORT-seq method (Muraro et al., 2016) (Muraro et al., 2016). Briefly, first and second strand synthesis (Invitrogen) was performed and all wells of a single plate were pooled. After in vitro transcription (Ambion), the amplified RNA was reverse transcribed and amplified for 10-12 cycles with Illumina Truseq primers. Finally, libraries were analyzed on an Illumina NextSeq500 using 75-bp pair-end sequencing.

## Analysis single-Cell mRNA Sequencing

Unique molecular identifier (UMI) count matrices were imported into R Studio and processed with the R package Seurat (version 3.1) (Stuart et al., 2019). For QC, we quantified the proportions of UMIs mapped to the mitochondrial genome. All the cells with mitochondrial reads > 10% were excluded. We further filtered cells with >7500 unique features in all 206 control and 216 ISX cells were passed to analysis. ERCC92 spike-ins as well as genes associated with clustering artefacts (Rn45s, Malat1, Kcnq1ot1, A630089N07Rik) were also excluded from the final dataset. Control and treated data sets were merged after QC filtering and then split by treatment for normalisation using the SCTransform wrapper and percent mitochondrial variations regressed. We calculated a subset of 3000 features to integrate using the SelectIntegrationFeatures and ensured all pearson residuals were calculated using PrepSCTIntegration. The data sets were then integrated using the FindIntegrationAnchors and the IntegrateData functions. We then ran an integrated analysis on all cells in the experiment using the standard workflow using default parameters: linear dimensional reduction using RunPCA, non-linear dimensional reduction using RunUMAP. A K-nearest neighbour’s graph was constructed with the FindNeighbors function using the first 25 principle components before clustering using the FindClusters function with a resolution of 1.8. Differential gene analysis for cluster definition was run using the FindConservedMarkers which reports the top markers conserved between the control and treated groups.

## Serotonin release from mouse and human small intestine organoids

Serotonin secretion in control and ISX-9 treated mouse organoids was measured after 7 days in culture and in human organoids after 10 days in culture. A 48-hr ISX-9 pulse was given to both mouse and human organoids as described above. At the end of the culture period organoids were released from BME and Matrigel with Cell Recovery solution (Corning), were washed with PBS and incubated with HEPES saline buffer (of 4.5 mM KCl, 138 mM NaCl, 4.2 mM NaHCO_3_, 1.2mM NaH_2_PO_4_.2H_2_O, 2.6mM CaCl_2_, 1.2mM MgCl_2_ and 10mM HEPES (pH7.4)) with 0.5% BSA for 2 hrs. Serotonin secretion was stimulated with 10 μM IBMX and 10 μM Forskolin in the presence of 1 μM fluoxetine (for blocking potential serotonin reuptake via SERT) for 1 hr. Supernatant and organoid lysates were collected for measuring secretion and content, respectively. Serotonin concentration was measured using an ELISA kit according to manufacturer’s instructions.

## Calcium imaging

30mm petri dishes were coated with BME diluted in Advanced DMEM/F12 (1:100) and left to set at 37 °C for 1 hr. Mouse small intestine organoids were collected and washed 3 times with cold PBS to dissolve BME and re-seeded in BME-coated petri dishes with ENR medium. After 2 hrs, organoids were loaded with 7μM Fura 2-AM in HEPES saline buffer containing 0.01% pluronic F127 and 2 mM probenecid, and incubated for 30 min at 37 °C. Organoids were washed 3 times with HEPES buffer and images were recorded with an Axiovert 135 Ca^2+^ imaging system (Zeiss), at 20x magnification every 80 milliseconds. Cells were excited at 340 nm and 380 nm and emitted light was acquired at 510 nm. Calcium concentration was calculated by 340/380 fluorescence ratio using a MetaFluor software. ATP (100μM) was used as positive control. Imaging experiments were performed at least 3 times and representative time course is presented in manuscript.

## Genetic engineering of human terminal ileal organoids

For overexpression of *Pax4* in human terminal ileal organoids, mouse *Pax4* cDNA was cloned into the pPB-tetO-MCS-IRES-mCherry vector. Briefly, pPB-tetO-MCS-IRES-mCherry vector was first linearized following sequential digestion with XhoI and SpeI restriction enzymes and gel purified using GenElute™ Gel Extraction Kit (Sigma). PCR primers for mouse *Pax4* were designed with 15 bp extensions that are complementary to the ends of the pPB-tetO-MCS-IRES-mCherry linearized vector. The 15bp extension were required for directional cloning using the In-Fusion HD cloning kit (Clonetech). From this point on, manufacturer instructions were followed. Pax4 was amplified using as template cDNA generated from mouse intestinal organoids. The generated plasmid was named pPB-tetO-*Pax4*-IRES-mCherry. The simultaneous co-transfection of 3 different plasmid is required for the generation of inducible overexpressing Pax4 human terminal ileal organoids. These plasmids are: pPB-tetO-*Pax4*-IRES-mCherry, pPB-CAG-rtTA-IRES-Hygro and pCAG-PBase plasmid. For electroporation of the 3 plasmids into human terminal ileal cells we followed the protocol from Fujii *et al (Fujii et al., 2015)*. DNA was transfected at 7.2 μg for the two piggyBac vectors, and at 5 μg for the transposase vector. 5 days post-electroporation cells were selected with 100 μg/ml hygromycin. Gene expression was induced using 1 μg/ml of Doxycycline.

## Statistics

For cell counting experiments, “n” represents the number of individual organoids assessed. For RNA experiments, “n” represents the number of biological replicates. All data are presented as mean ± standard error of the mean (SEM), except violin plots in which data is presented as median and quartiles. Each “n” is presented as a dot in graphs. Relevant tests described in figure legends. All statistical analyses were performed using Graph Pad Prism Version 8.1.2 (GraphPad Software) for Windows or Mac except for RNA-seq where statistics were calculated using Seurat.

