## Supplementary material for "ISX-9 manipulates endocrine progenitor fate revealing conserved intestinal lineages in mouse and human": All supplemental figures

### Supplementary Figure 1

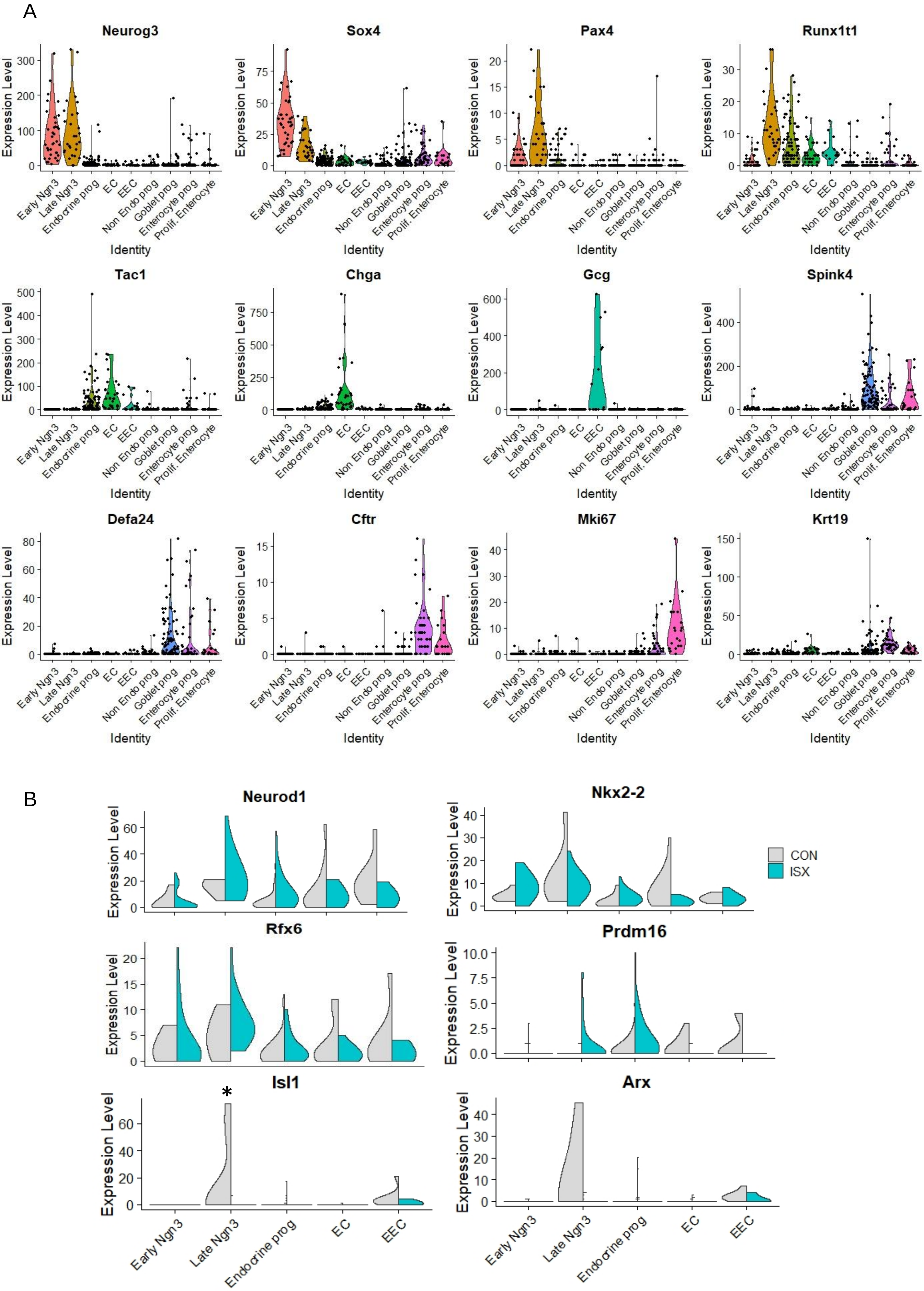

### Supplementary Figure 2

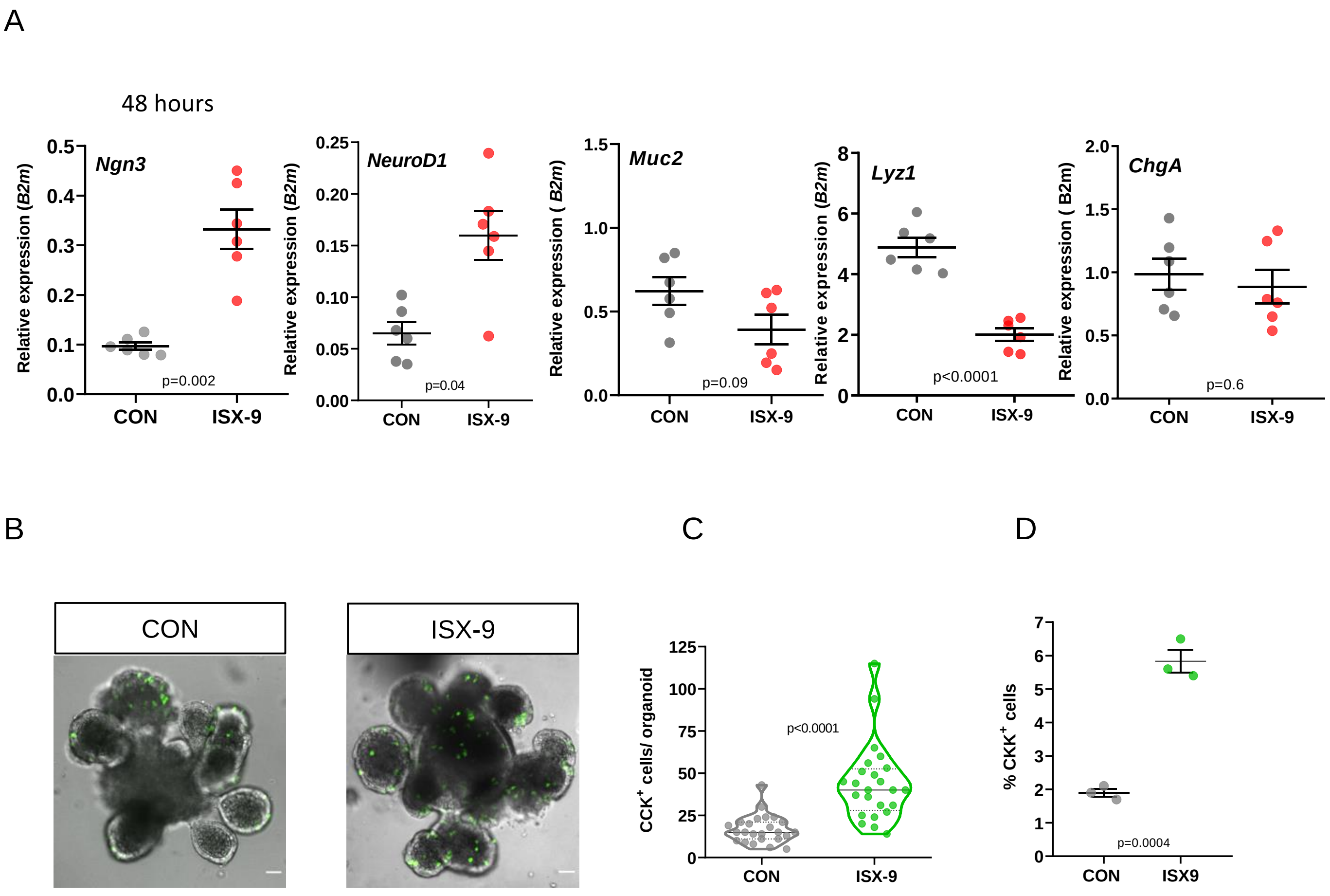

### Supplementary Figure 3

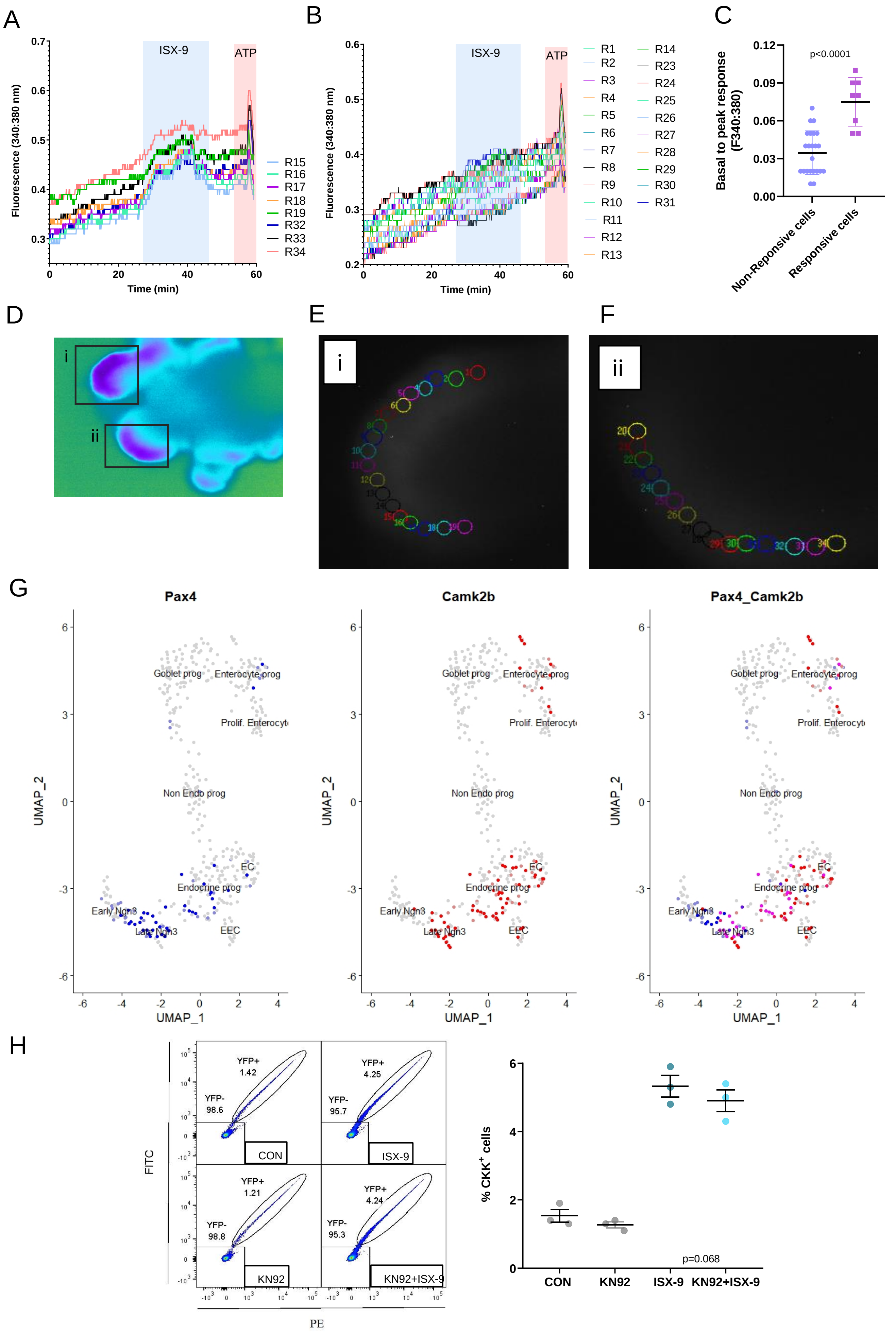

### Supplementary Figure 4

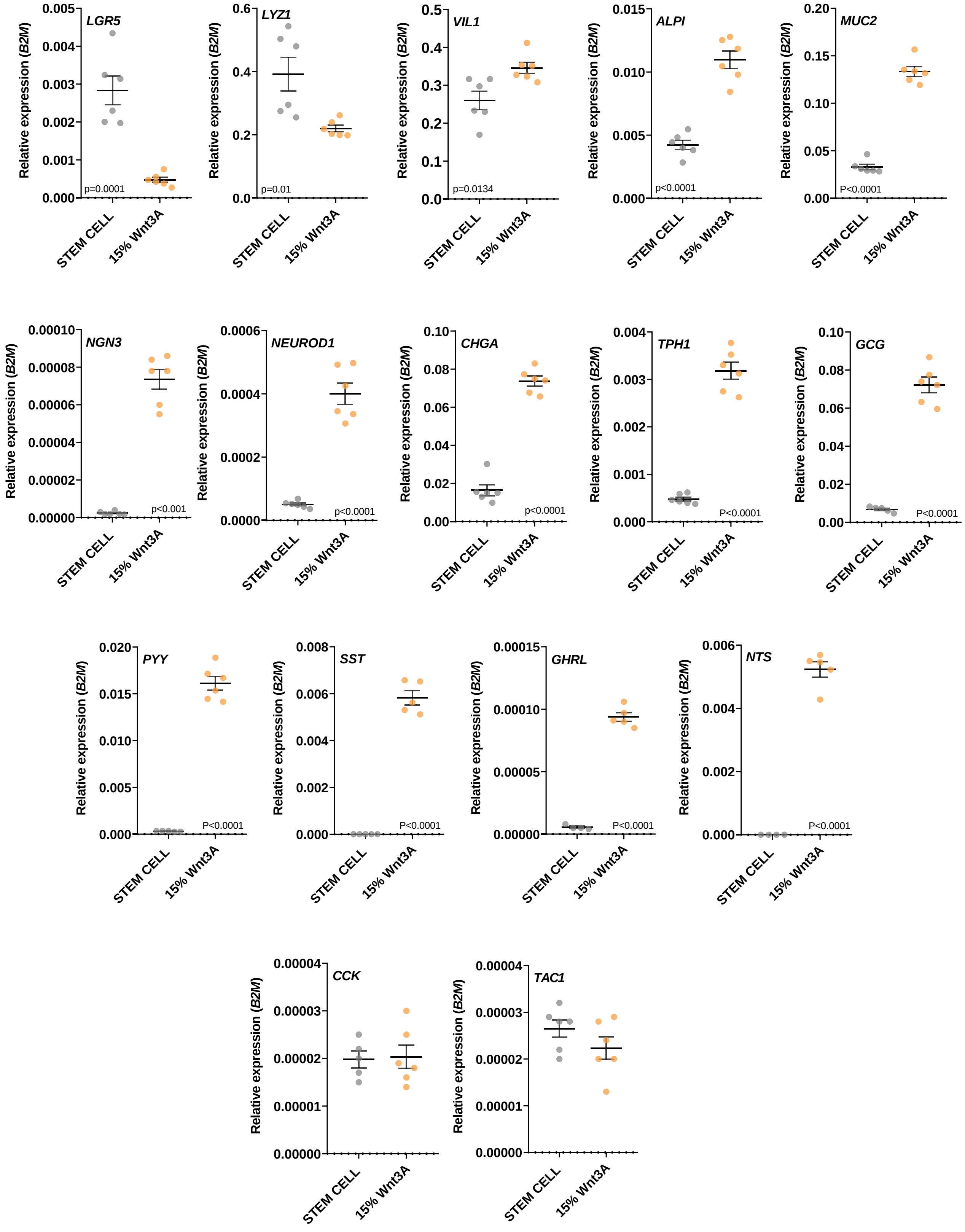

### Supplementary Figure 5

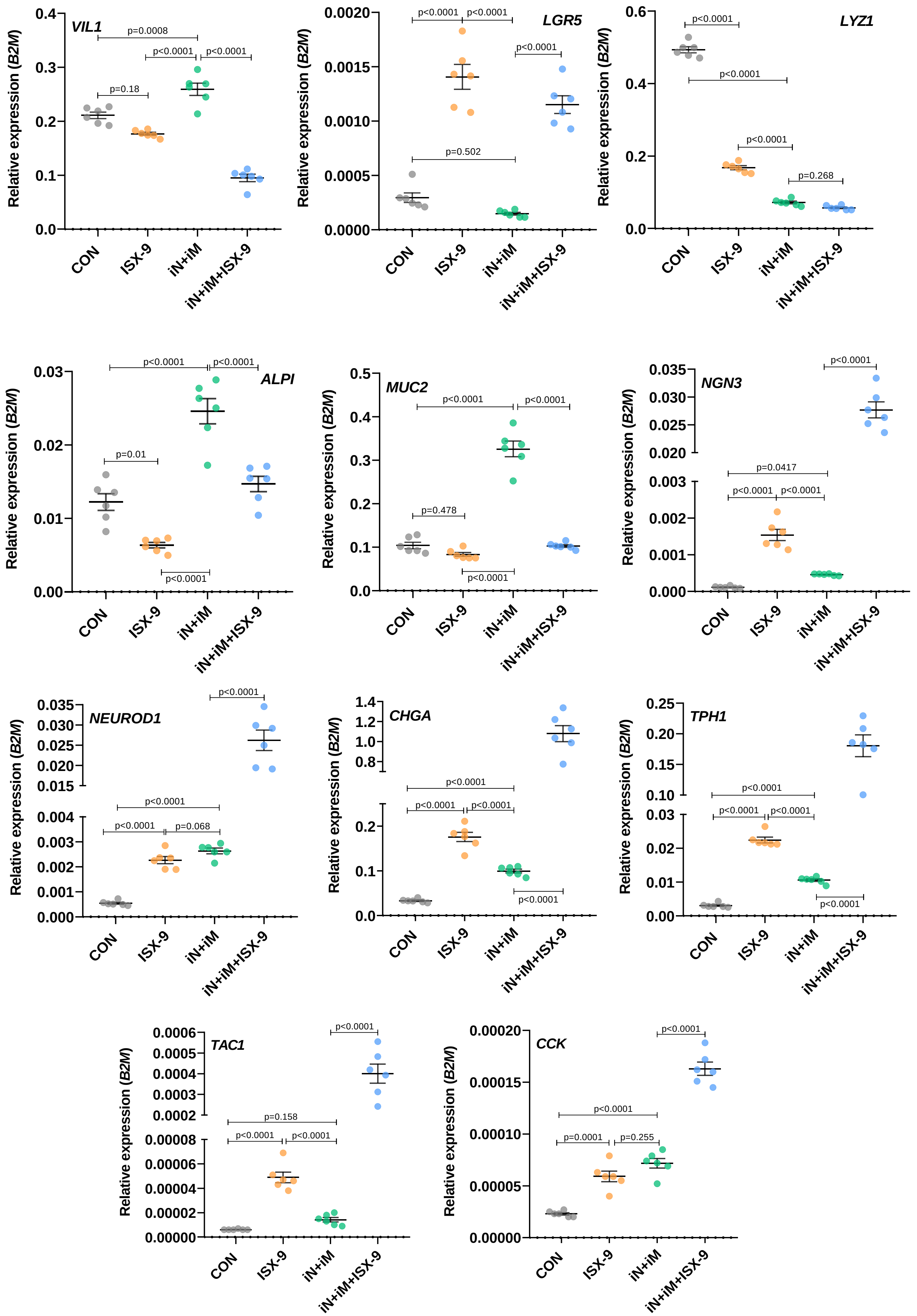

### Supplementary Figure 6

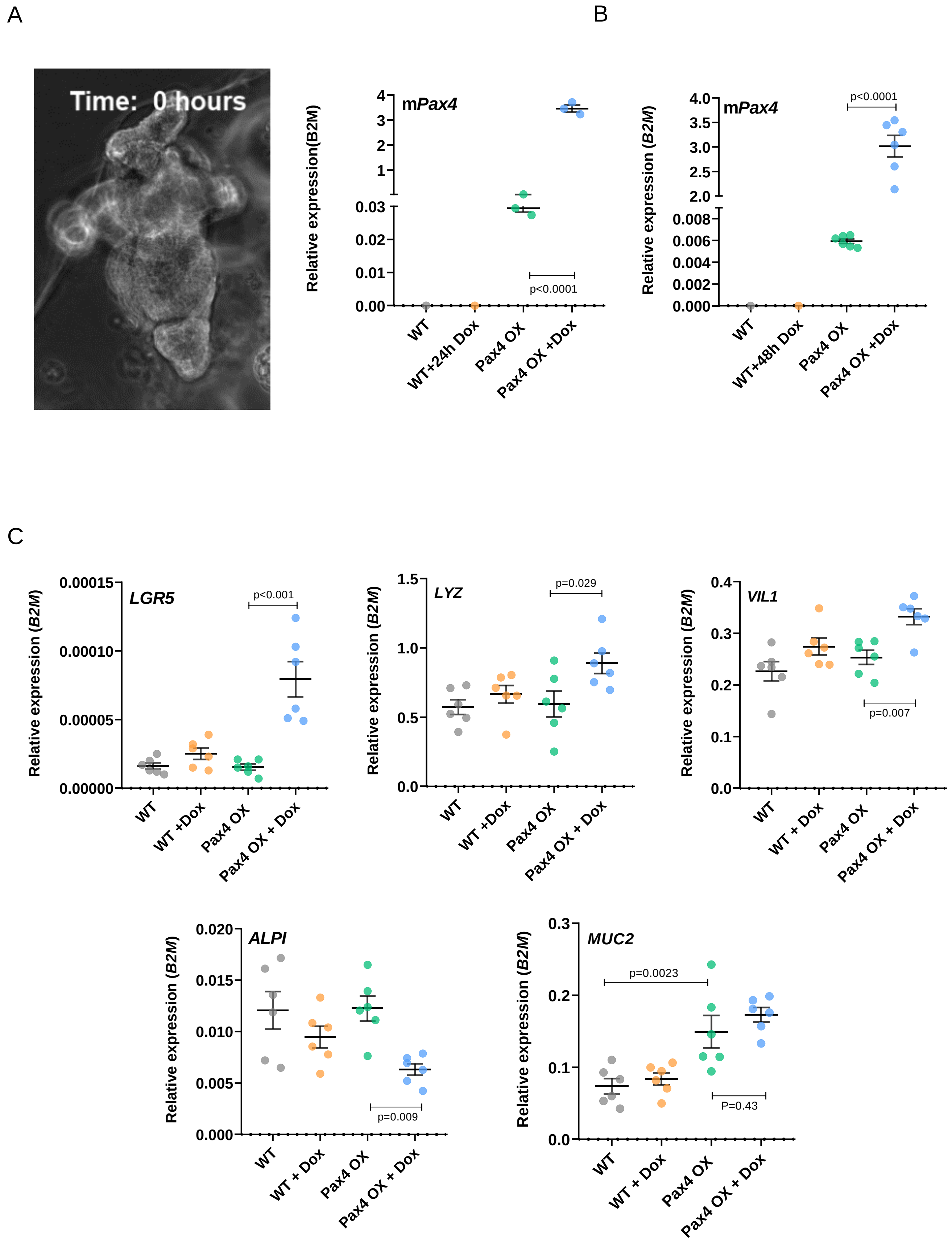

### Supplementary Figure 7

A

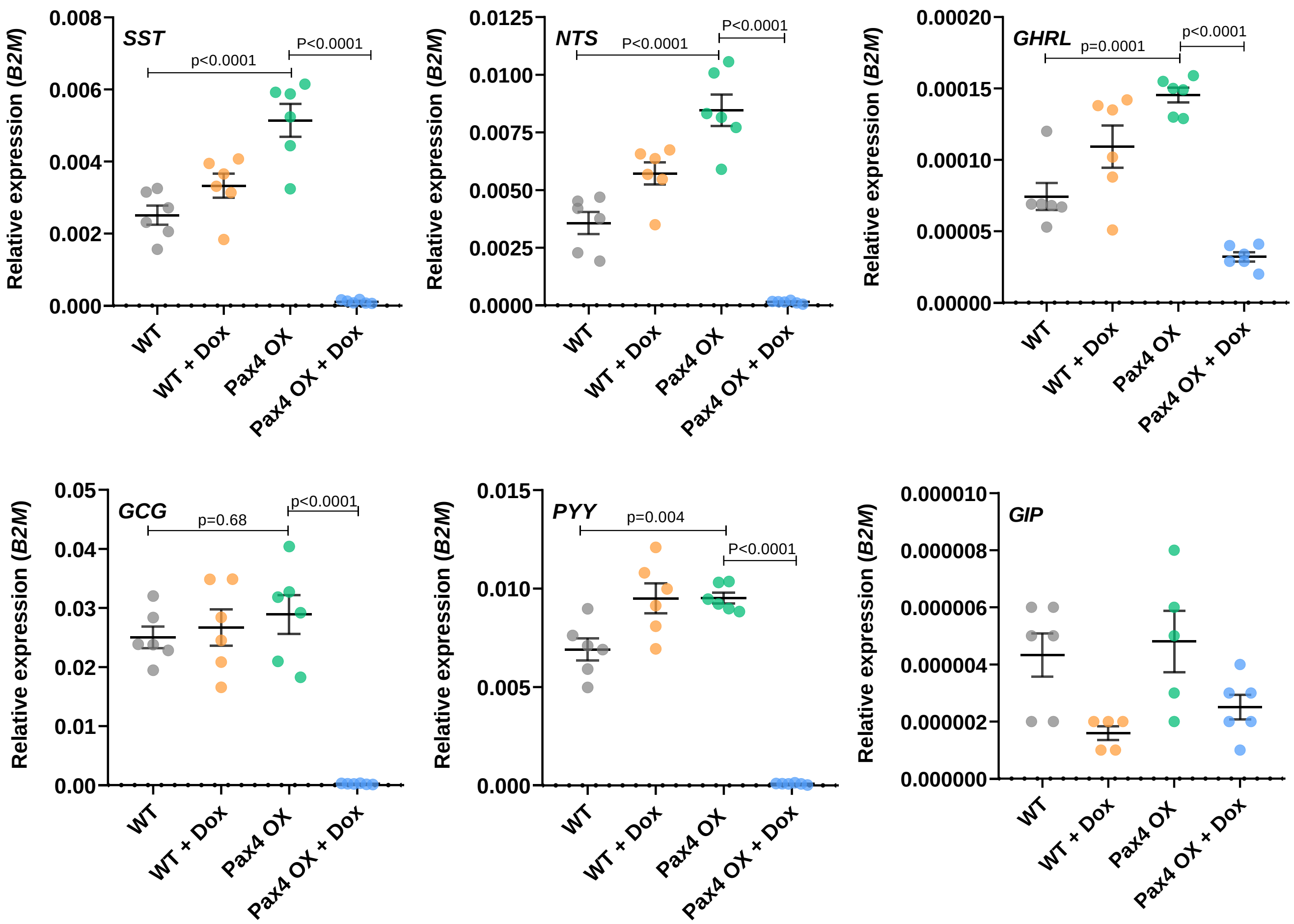


B

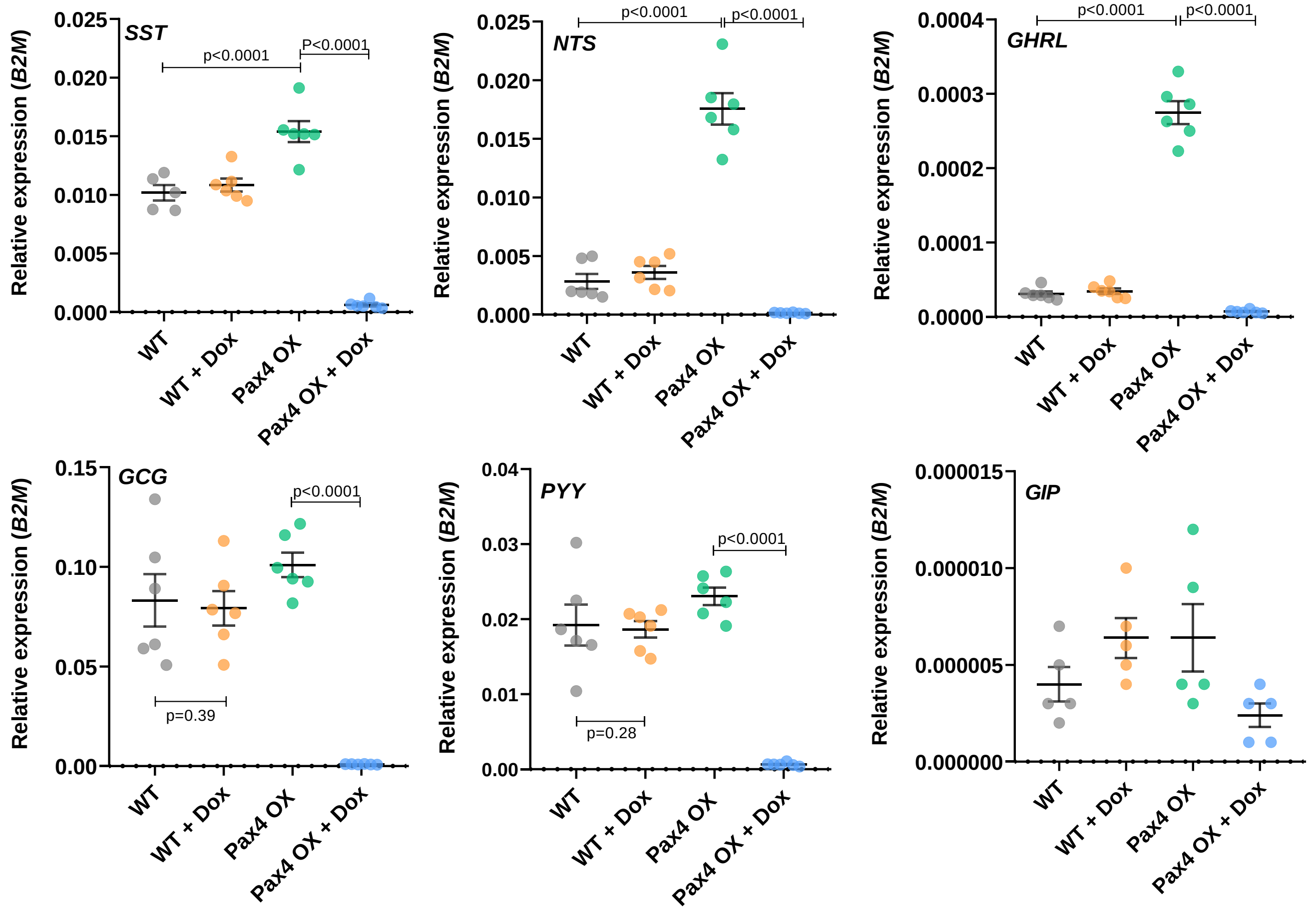
